# Heritability of movie-evoked brain activity and connectivity

**DOI:** 10.1101/2024.09.16.612469

**Authors:** David C. Gruskin, Daniel J. Vieira, Jessica K. Lee, Gaurav H. Patel

## Abstract

The neural bases of sensory processing are conserved across people, but no two individuals experience the same stimulus in exactly the same way. Recent work has established that the idiosyncratic nature of subjective experience is underpinned by individual variability in brain responses to sensory information. However, the fundamental origins of this individual variability have yet to be systematically investigated. Here, we establish a genetic basis for individual differences in sensory processing by quantifying (1) the heritability of high-dimensional brain responses to movies and (2) the extent to which this heritability is grounded in lower-level aspects of brain function. Specifically, we leverage 7T fMRI data collected from a twin sample to first show that movie-evoked brain activity is heritable across the cortex, and that this heritability is greater for information encoded in lower temporal frequencies, especially in more associative cortical areas. Next, we use hyperalignment to decompose this heritability into genetic similarity in *where* vs. *how* sensory information is processed. We also show that the heritability of brain activity patterns can be partially explained by the heritability of the neural timescale, a one-dimensional measure of local circuit functioning. Finally, we generalize our findings by illustrating a similar pattern of results for the heritability of movie-evoked functional connectivity. These results demonstrate that brain responses to complex stimuli are heritable, and that this heritability is due, in part, to genetic control over stable aspects of brain function.

## Introduction

Although the neural machinery that allows us to process sensory information is broadly conserved across people, no two individuals experience the same sensory stimulus in exactly the same way. What underlying factors give rise to this person-to-person variability in subjective experience? Recent work has established that the idiosyncratic nature of subjective experience is reflected in idiosyncratic brain responses to sensory information, and that how an individual processes a stimulus is shaped by their psychosocial background and previous experiences. For example, individuals who share more similar personality traits (Finn et al., 2018) and political orientations (van Baar et al., 2021) exhibit more similar interpretations of, and functional magnetic resonance imaging (fMRI) responses to, relevant audiovisual stimuli, as do individuals who are primed with more similar contextual information before listening to an ambiguous narrative (Yeshurun et al., 2017). Here, we extend this work by investigating a more fundamental source of variability in sensory-evoked brain responses and the experiences they represent: our genes.

Whether individual variability in a given trait is due to environmental or genetic factors is a central question in biology, and the extent to which this variability is underpinned by variation in genetics is captured by heritability (or *h*^2^). Recent studies have quantified the heritability of various aspects of sensory brain function, revealing a genetic basis for patterns of brain activity elicited by auditory tones and visual gratings in sensory cortices (Alvarez et al., 2021; Renvall et al., 2012; van Pelt et al., 2012). However, the unimodal and low-dimensional nature of these stimuli may not capture the full complexity of real-life sensory experiences and the brain responses they evoke. Consequently, the extent to which genetic factors influence brain responses to more naturalistic stimuli remains unclear, especially for high-level (e.g., social and narrative) information encoded across longer timescales in association cortex.

In addition to activity patterns within individual brain areas, information can also be encoded in the functional connectivity (FC) between multiple areas or networks (Chen et al., 2014; Kohn et al., 2016). Research into the heritability of FC and related measures has largely focused on data acquired while subjects are at rest, during which an individual’s unique FC profile (i.e., pattern of pairwise FC strengths), describes their brain’s intrinsic functional architecture (Anderson et al., 2021; Burger et al., 2022; Busch et al., 2023; Dworetsky et al., 2024; Glahn et al., 2010; Sinclair et al., 2015; van den Heuvel & Hulshoff Pol, 2010). These studies have demonstrated that a range of resting state FC (rest FC)-derived measures are moderately heritable, and similar findings have resulted from work characterizing the heritability of FC during task performance, which additionally reflects the processing of information relevant to the task at hand (Cole et al., 2021; Elliott et al., 2019; Korgaonkar et al., 2014). Although this work has shed significant light on the genetic basis of FC during rest and cognition, the heritability of sensory-evoked FC patterns, which are known to encode stimulus features (Chen et al., 2014) and track individual differences in behavior (Finn & Bandettini, 2021), has yet to be investigated.

Finally, brain activity and connectivity patterns are complex phenomena that arise from a variety of physiological processes. Although the heritability estimates established by previous work could reflect emergent aspects of brain function, it might instead be possible to reduce them to genetic control over lower-level neural, vascular, and metabolic processes. For example, although the human cortex is topographically organized into areas that are specialized for processing specific kinds of information (e.g., facial features), the locations of these areas and the tuning patterns within them vary widely across individuals (Gordon et al., 2017; Haxby et al., 2020; Petersen et al., 2024). As such, these cortical topographies, or individual-specific maps of *where* stimulus features are processed, emerge over the course of development and constrain the activity and connectivity patterns an individual will exhibit during sensory processing. Independent of *where* stimuli are processed, stable aspects of brain function also shape high-dimensional activity and connectivity patterns by influencing *how* information is processed. For example, recent work from Shinn et al. (2023) showed that individual variability in higher-order aspects of brain function like FC profiles can be traced back to variability in simpler, low-level phenomena like temporal autocorrelation of the blood oxygen level-dependent (BOLD) signal. More specifically, a measure of temporal autocorrelation known as the neural timescale (NT) is thought to reflect the strength of local recurrent excitation (Cavanagh et al., 2020) but is also closely tied to the organization of brain-wide FC profiles (Shinn et al., 2023). Given that lower-level properties like functional topography (Alvarez et al., 2021; Anderson et al., 2021; Dworetsky et al., 2024) and BOLD temporal autocorrelation (Christova et al., 2022) are themselves heritable, it remains unclear (1) to what extent high-dimensional brain responses to naturalistic stimuli are heritable and (2) how much of this heritability can be reduced to genetic control over these stable spatial and temporal aspects of brain function.

In the present work, we address these questions by analyzing 7T fMRI recordings of a twin sample acquired by the Human Connectome Project (Van Essen et al., 2013) to quantify the heritability of two distinct high-dimensional traits—stimulus-evoked BOLD time courses and functional connectivity profiles—across the cortex. Here, we focus on fMRI data acquired during movie-watching, as the rich and multimodal nature of movies engages multiple sensory and associative regions as well as the connections between them, making them well-suited for broadly assessing individual differences in sensory processing. Leveraging a multi-dimensional estimator of heritability (Anderson et al., 2021), we first show that movie-evoked BOLD time courses are heritable across the cortex. We extend this result by showing that BOLD time course heritability is greater in slower frequency bands, and especially in more associative parcels, suggesting that the neural processing of more abstract vs. lower-level sensory information is under greater genetic control. Next, we use hyperalignment to separate heritable differences in *where* information is processed from *how* it is processed by projecting voxel-level data into a common functional space, parsing the heritability of high-dimensional BOLD time courses into genetic control over stable spatial (e.g., functional cortical topography) versus temporal (e.g., neural timescale) aspects of brain function. Finally, we reveal a similar pattern of results for a different set of high-dimensional brain responses: functional connectivity profiles. Taken together, these results characterize the degree to which sensory processing is controlled by genetics and illustrate the benefits of a reductionist approach to studying the heritability of complex neurobiological phenomena, providing a foundation for future multi-scale studies of the mechanisms that underlie heritable differences in brain function.

## Methods

### Participants

Data used for this project come from the 178 subjects in the Human Connectome Project (HCP) Young Adult 7T release who completed every movie-watching run (Van Essen et al., 2013). All participants were healthy individuals between the ages of 22 and 36 (mean age = 29.4 years, standard deviation = 3.3) and provided informed written consent as part of their participation in the study. Self-reported racial identity in this sample was 87.6% White, 7.3% Black or African American, 3.9% Asian/Native Hawaiian/Other Pacific Islander, and 1.1% unknown/not reported, and 1.7% of the sample identified as Hispanic/Latino. HCP twin zygosity was determined by genotyping (168 subjects) or self-report (4 subjects), which identified 51 monozygotic (MZ) twin pairs and 34 dizygotic (DZ) twin pairs in the present sample, as well as 2 pairs of non-twin siblings and 4 singletons. All sibling pairs shared the same gender. Out of these 178 subjects, we identified 690 unrelated dyads who were matched in gender and age in years. Because two of these participants (from two separate MZ twin pairs) did not complete every resting state run, analyses involving resting state data use a sample size of *n* = 176.

### fMRI data

All fMRI data were collected on a 7T Siemens Magnetom scanner across four sessions spanning multiple days. Each day involved two resting state (900 volumes) and two movie-watching scans (variable durations) across two sessions, all with the following sequence: repetition time (TR) = 1000 ms, echo time (TE) = 22.2 ms, number of slices = 85, flip angle = 45 degrees, spatial resolution = 1.6 mm^3^. During movie runs, subjects passively watched short audiovisual clips from either independent films or major motion pictures as well as a montage of brief videos. All clips were only viewed once by each subject, with the exception of the brief montage which was included at the end of each of the four runs for test-retest purposes. These clips differed in their degree of narrative and social content, but many featured language and human characters; more information on the clips can be found at https://db.humanconnectome.org. Each video clip was preceded by 20 seconds of rest, so we discarded all volumes that took place during these rest blocks as well as the first 20 volumes of each clip to prevent rest data and onset transients from biasing our intersubject correlation (ISC) measurements. Rest and movie data from the same day were normalized and concatenated, yielding one rest run (1800 volumes) and one movie run (1432 volumes Day 1, 1409 volumes Day 2) for each day of data collection.

### Preprocessing and parcellation

The fMRI data used here were preprocessed as described in a previous publication (Gruskin & Patel, 2022). Briefly, we used ICA-FIX denoised data from which the global signal and its temporal derivative were removed. Our use of global signal regression (GSR) was motivated by work showing that this approach effectively reduces the impact of nuisance signals on FC measures (Parkes et al., 2018) and increases relationships between measures of brain function and behavior (Li et al., 2019) across individuals (presumably by making individual FC profiles more distinguishable), both of which should highlight heritable aspects of brain function. Still, it is important to consider that GSR remains a controversial preprocessing step and may affect our results. As such, we repeated our main BOLD time course heritability analysis without using GSR. To examine the effects of parcellation resolution, data were parcellated using the 10 resolutions of the Schaefer atlas (100 to 1000 parcels; Schaefer et al., 2018). The ICA-FIX data downloaded here were aligned across subjects using the HCP’s Multimodal Surface Matching (MSMAll, henceforth “MSM”) method, which registers data based on several multimodal properties in a topology-preserving manner (Feilong et al., 2021; Robinson et al., 2014).

### Intersubject correlation (ISC)

Dyadic ISC analyses were used to quantify BOLD time course similarity between all pairs of participants. For each vertex or parcel, each participant’s BOLD signal time course from a given day’s movie-watching scan was normalized and (Pearson) correlated with the corresponding BOLD signal time courses from all other participants to yield an ISC matrix. We chose to use this pairwise ISC method over a leave-one-subject-out approach because it allows us to capitalize on the information contained in the *n*^2^ pairwise ISC matrix (whereas the other approach averages out meaningful information to yield an *n* × 1 ISC matrix). All Pearson *r* values in this and all other analyses were Fisher *z*-transformed before averaging (and converted back to Pearson *r* for visualization). This analysis was intended to illustrate group-level dyadic BOLD time course similarity in familiar units, not to test for significant group difference effects, as individuals may have contributed to multiple dyads such that each dyad was not independent. As such, we reserved hypothesis testing for our formal heritability analyses (below).

### Functional connectivity (FC)

We constructed resting state functional connectivity (rest FC) and movie-watching functional connectivity (movie FC) matrices by (Pearson) correlating the time courses of all parcel pairings from the 400-parcel Schaefer atlas, using data from the two concatenated rest scans. These matrices were produced for each subject and for each day of data collection. We applied the same procedure to the movie-watching data, enabling us to compare the heritability of rest and movie FC profile similarities and strengths.

To evaluate the similarity of FC profiles for each combination of the 17 networks defined by Kong et al. (2019), we first vectorized the FC matrices of each subject. We then extracted the correlation coefficients corresponding to the parcel-level connections comprising each Kong network combination and computed correlations for these vectorized profiles across all subject pairs. For example, there are 16 parcels in the Kong et al. Auditory network and 17 parcels in the Language network, so the FC profile for a given subject’s Auditory-Language network combination consists of the 272 (16*Auditoryparcels* × 17*Languageparcels*) correlation coefficients between all unique pairs of one parcel from each network. We also assessed subject-level FC strengths for each network combination by averaging the correlation coefficients for all parcel-level connections within a network combination. As above, this analysis was intended to illustrate twin-twin similarity in familiar units, and given dyadic interdependence issues we reserved hypothesis testing for our formal heritability analyses (next section).

### ISC and FC profile heritability analyses

BOLD time courses and FC profiles are high-dimensional variables, and reducing their dimensionality in order to use classical heritability analyses would sacrifice both statistical power and interpretability. As such, we quantified their heritability with a multidimensional estimator that has been used in similar studies (Anderson et al., 2021; Busch et al., 2023; Ge et al., 2016). This model (detailed in Anderson et al., 2021) takes as input a Subjects × Subjects kinship matrix describing the degree of genetic relatedness between individuals (1 for MZ twins, 0.5 for non-MZ siblings, 0 for all other pairs) as well as a Subjects × Subjects phenotypic similarity matrix. Here, each value of the phenotypic similarity matrix corresponded to a Pearson correlation coefficient describing either BOLD time course (per parcel or voxel) or FC profile similarity (per network combination) for a given subject pair. To estimate heritability, the variance in phenotypic similarity is then partitioned into a component attributable to genetic factors, represented by the kinship matrix, with age, gender, and per-scan head motion included as covariates. Significance testing of individual multidimensional heritability values and calculation of their standard errors were performed using the method established by Anderson et al. 2021. Specifically, the kinship matrix was shuffled 10,000 times to generate a null distribution against which the observed value could be compared. The resulting p-values were then false discovery rate (FDR) corrected using the Benjamini-Hochberg method (Benjamini & Hochberg, 1995) to adjust for multiple comparisons. FDR correction was always performed separately for each day of data collection, such that results reported as “significant at FDR-corrected *P* < .05 on both days” reflect a conservative criteria of *q* < .05 on two independent tests. Standard errors (SEs) were derived through a block jackknife method in which heritability was recalculated 90 times after leaving out all members of one of the 90 families in the dataset on each iteration. We then used these SEs to generate 95% confidence intervals (CIs).

To compare the heritability of movie and rest FC profiles, we used the following non-parametric permutation approach. For each day of data collection, we randomly shuffled each subject’s movie and rest FC matrices (along with their corresponding framewise displacement [FD] covariates) and re-calculated FC profile heritability using the shuffled FC matrices. We then subtracted the two resulting heritability values for each of the 153 unique network combinations for the 17 Kong networks (e.g., Auditory-Language, Auditory-Auditory, etc.) and averaged these across networks to obtain 17 values reflecting the null difference in FC profile heritability for each network. We repeated this procedure 10,000 times and then used the two-sided test described above to generate a p-value for each network, and these p-values were then FDR corrected separately for each day of data collection. Because complete resting state datasets were not available for 2/178 movie-watching subjects, we only used movie-watching data from the 176 subjects who also had complete resting state data for this permutation test.

To determine the sample size necessary for stable multidimensional heritability results, we conducted our BOLD time course heritability analysis multiple times while systematically excluding between 5% and 90% of families. At each exclusion level, we performed 100 iterations, each time randomly removing a subset of families. After each iteration, we calculated the absolute difference between the heritability values obtained from the subsample and those from the full sample for each parcel. We then averaged these differences across all parcels and iterations to obtain the mean absolute error for that exclusion level. Similarly, to assess the stability of the spatial pattern of our results, we computed Spearman correlations between the subsample and full sample heritability values across parcels and averaged these correlations across all iterations. We included this analysis because it serves as a simple way to demonstrate the stability of our results at various sample sizes, and because this sort of subsampling approach has been used many times before in our field (e.g., Marek et al., 2022) and others (e.g., Manyara et al., 2024) to demonstrate the sample-size dependence of statistical effects.

Because the heritability of ISC is constrained by the degree of synchronization in a given area, we also sought to identify areas in which BOLD time courses were more/less heritable than would be expected based on ISC alone by fitting a linear model of the form heritability_*i*_ = *β*_0_ + *β*_1_ ⋅ ISC_*i*_ + *ε*_*i*_ and plotting the residuals.

### Frequency-dependent ISC heritability analyses

To characterize the frequency-specific heritability of movie-evoked BOLD time courses, we performed a spectral analysis of the BOLD time series data across all subjects and parcels. First, we set a high-frequency cutoff equal to the Nyquist frequency of our data (0.5 Hz) and defined a low-frequency cutoff at 0.004 (1/238) Hz, corresponding to the length of the longest clip. We then computed the power spectrum for each parcel, subject, and day of data collection by applying a Fast Fourier Transform (FFT) to the time series data of each subject and then averaging these spectra across all subjects, parcels, and scanning days. To ensure consistent comparison across frequencies, we interpolated the power spectra onto a common frequency axis with a resolution of 0.001 Hz. The cumulative power distribution was then calculated from this averaged power spectrum. To partition the frequency range into bands containing equal fractions of the total power, we identified frequency cutoffs corresponding to quintiles of the cumulative power distribution. This resulted in five frequency bands: Band 5 (0.004–0.02 Hz), Band 4 (0.02–0.04 Hz), Band 3 (0.04–0.07 Hz), Band 2 (0.07–0.14 Hz), and Band 1 (0.14–0.50 Hz). For each of these bands, we applied fourth-order Butterworth bandpass filters to the concatenated BOLD time courses of each subject and parcel and recalculated ISC and BOLD time course heritability as described above. Standard errors of cortex-wide heritability were estimated for each band using a leave-one-family-out jackknife procedure. Heritability estimates were first averaged across all brain parcels within each jackknife fold, and the standard error of these parcel-averaged values was computed using the standard jackknife variance formula. To further characterize frequency-dependent changes in heritability at the parcel level, we then Spearman-correlated the spatial patterns of heritability with the sensorimotor-association hierarchy rankings from Sydnor et al. (2023). Significance testing for these sensorimotor-association results was performed using BrainSMASH (Brain Surrogate Maps with Autocorrelated Spatial Heterogeneity; Burt et al., 2020).

### FC strength heritability analysis

Because FC strengths are inherently one-dimensional traits, their heritability was quantified with SOLAR’s *polygenic* function (Almasy & Blangero, 1998), which generates estimates using variance-component models. Age, gender, and head motion were used as covariates in all FC strength analyses, and SEs were calculated using the block jackknife procedure described above. We note that because the heritability of univariate (vs. multivariate) traits requires larger sample sizes, these results should be considered preliminary and would benefit from further investigation in larger samples (Anderson et al., 2021). To test the significance of differences in rest vs. movie FC strength heritability, we compared the observed differences to null distributions generated by 1,000 permutations of the same procedure described above for FC profiles.

### Hyperalignment

We used piecewise response and connectivity hyperalignment (RHA and CHA, respectively), two complementary methods for aligning data into a topography-independent common space, to functionally align vertex-level BOLD time courses across subjects (Guntupalli et al., 2018; Haxby et al., 2011). We decided to use piecewise hyperalignment, in which vertices are aligned within non-overlapping parcels, instead of searchlight hyperalignment because it has been shown to be both more accurate and more efficient (Bazeille et al., 2021). This approach independently transforms each subject’s data within discrete anatomical parcels into the common space, yielding functionally aligned vertex time series that are calculated as weighted linear combinations of the original time series from all other vertices within that same parcel for that subject. We then repeated our BOLD time course and FC profile heritability analyses using these hyperaligned datasets to quantify the extent to which brain response heritability reflects genetic control over cortical topography. *RHA:* For each Schaefer atlas parcel and day of data collection, we used iterative Procrustes transformations to align vertex-level BOLD time courses to a common model information space. This yielded one invertible transformation matrix per parcel and per subject, which we then used to project data from the other day of data collection (which was not used to generate the transformation matrices) into the common information space. *CHA:* The same iterative Procrustes approach was used for CHA, but here the input data consisted of rest FC profiles for each vertex within a given parcel. Each vertex’s functional connectivity profile consisted of the Pearson correlation coefficients between that vertex’s time course and the average time courses from all other parcels, the number of which varied with different parcellation resolutions. After training a model whose dimensions correspond to shared rest FC properties (instead of the shared response properties in RHA), we used the corresponding transformation matrices to align movie-watching time series data from the other day of data collection. To quantify the spatial scale at which cortical topographies contribute to brain response heritability, we repeated the RHA and CHA procedures described above for each of the 10 Schaefer atlas resolutions to yield 21 datasets per subject (10 RHA-aligned datasets, 10 CHA-aligned datasets, and the original MSM-aligned dataset).

### Relationships between parcel area and heritability

We used power law modeling to characterize the relationship between cortex-level heritability and hyperalignment area. To calculate parcel areas, we first generated vertex-level areas using the *-surface-vertex-areas* function in wb_command and then summed the areas of all vertices included in each parcel. Next, for each hyperalignment method and each day of data collection, we used nonlinear least squares regression to fit a power law model (*y* = *a* ⋅ *x*^*b*^ + *c*) to the 11 heritability values calculated from the 10 Schaefer atlas resolutions and one from MSM-only aligned data and the 10 average Schaefer atlas parcel areas as well as 0 (corresponding to no hyperalignment in the MSM-only data).

### Neural timescale (NT) analyses

To determine the extent to which BOLD time course heritability reflects genetic control over NT, we took an approach used in numerous studies to calculate NT at rest (known as intrinsic neural timescale, or INT; Watanabe et al., 2019; Wengler et al., 2020) and applied it to movie-watching data. NTs were calculated separately for each day of data collection as the sum of the autocorrelation coefficients from the first lag until the first lagged timepoint with a non-positive autocorrelation coefficient (Wengler et al., 2020) for each vertex.

Because time courses that are themselves more temporally autocorrelated will have a higher variance in their correlations with each other (Shinn et al., 2023), and because stimulus-evoked BOLD time courses tend to be positively correlated across subjects, we reasoned that pairs of individuals with longer collective NTs would have more correlated BOLD time courses. To test this directly, we (Spearman) correlated ISC values from one day of data collection with NTs calculated using the other day’s data across all possible subject pairs. Because our hypothesis here was unrelated to genetic effects, we tested the significance of the resulting vertex-wise correlation coefficients using a family-compliant quadratic assignment procedure with 1,000 permutations. To account for familial dependencies, we shuffled family units before shuffling individuals within those families. Two-tailed p-values were defined as the proportion of null correlations exceeding the observed coefficient. Finally, we applied FDR correction across all vertices for each day separately, considering results significant only if they were consistent across both independent scan sessions.

We then averaged NT values across all vertices to get a single, cortex-wide NT measure for each subject and each day of data collection. Dyadic NT similarity was quantified as the absolute value of the difference between each subject pair’s cortex-wide NT values. This differencing approach yielded several extreme values. We thus used the 1.5×IQR method to identify 4% of dyadic NT values as outliers and excluded them from the group differences analyses. Because these dyads are not independent (as discussed above), we used SOLAR to quantify the heritability of whole-brain NT (averaged across all vertices) and test for statistical significance, controlling for age, gender, and head motion. We also used the aforementioned multidimensional heritability analysis to quantify the heritability of parcel-level NT topographies (where the phenotypic similarity matrix reflects the Spearman correlation of each pair’s 400 × 1 parcel-level NT values).

We also included subject-level NT as a covariate in some multidimensional heritability analyses. When calculating heritability at a given vertex for one day of data collection, the NTs for that vertex from the other day of data collection were used as covariates. To evaluate whether including NT as a covariate decreased BOLD time course heritability, we generated a null distribution of 1,000 heritability values by randomly shuffling vertex-level NT vectors such that the NTs for one subject were paired with the ISC values and covariates from another subject. We then calculated two-sided permutation p-values.

### Significance testing for autocorrelated brain maps and FC matrices

We used Spearman correlations to quantify the reliability of ISC and FC heritability maps across days, as well as relationships between these heritability maps and the sensorimotor-association hierarchy ranking from Sydnor et al. (2023). Because cortical spatial maps are significantly autocorrelated, we used the variogram matching approach from BrainSMASH to assess the significance of these correlations. Briefly, this test works by generating autocorrelation-matched surrogates for one of the empirical maps from each correlation, calculating Spearman correlations between these surrogates and the other empirical map, and then comparing these null correlations to the correlation between both empirical maps. We performed independent tests using 1,000 unique surrogates for each hemisphere and averaged the two p-values to get the whole-cortex p-values we report in this manuscript. Because values in FC heritability matrices are similarly not independent of each other, we used Mantel tests with 10,000 permutations to test their reliability.

### Data and code availability

The raw HCP data used for this project can be downloaded from ConnectomeDB (db.humanconnectome. Code for all analyses will be posted on GitHub upon publication. Analyses were performed in MATLAB (R2023b), Python, and R. All cortical surface visualizations were performed with Connectome Workbench (Marcus et al., 2011).

## Results

### Similarity in movie-evoked brain activity increases with genetic relatedness

To characterize the heritability of brain responses to complex stimuli, we used 7T fMRI data from 178 HCP Young Adult subjects acquired across two days (using two largely non-overlapping sets of movie stimuli, see Methods) to (1) quantify the heritability of brain activity and connectivity patterns during movie-watching and (2) determine the extent to which the heritability of these dynamic, high-dimensional brain responses is grounded in stable and fundamental aspects of brain function like cortical topographies and neural timescales.

We first aimed to determine how closely movie-evoked brain responses were shared among pairs of individuals, and whether this similarity was influenced by their genetic relationship. Using inter-subject correlation (ISC) of parcellated BOLD time courses to index brain activity similarity, we found that identical (or monozygotic, MZ), fraternal (or dizygotic, DZ) and age- and gender-matched unrelated (UR) dyads differed in their level of brain activity similarity in a manner consistent with their relative degrees of genetic relatedness (although spatial distributions of ISC were consistent across groups, Fig. S1). More specifically, identical twins’ BOLD time courses were 59% more similar than those from pairs of unrelated individuals. When comparing identical twins to fraternal twins, the identical twins’ brain activity was still more similar, but by a smaller margin of 24%. Finally, fraternal twins had 29% more similar time courses than unrelated pairs. We observed the greatest group differences in parcels with medium levels of ISC (separation between the three traces in Fig. 1B, visualized as scatterplots in Fig. S2), suggesting that floor and ceiling effects may limit the degree to which genetic relatedness impacts brain activity in regions that are not driven by audio-visual stimuli and regions that exhibit highly stereotyped activity across all subjects, respectively.

**Figure 1.**
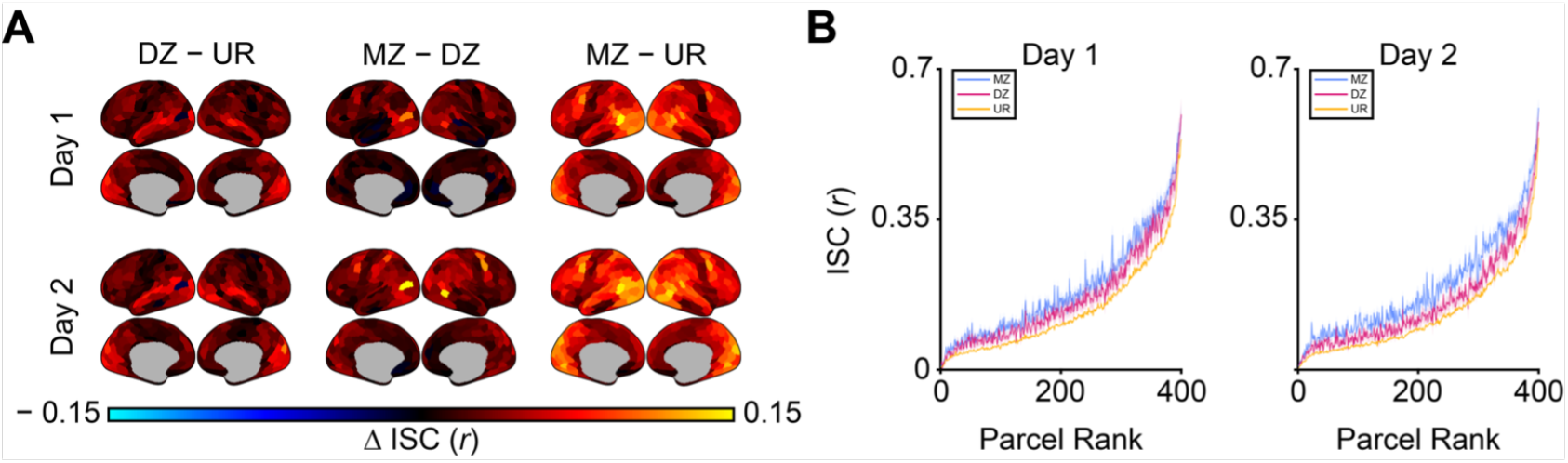
BOLD time course similarity scales with genetic relatedness across the cortex. (A) Group differences in average BOLD time course similarity (indexed by ISC) show that BOLD time course similarity is greater among dyads who are more genetically related (51 MZ dyads, 34 DZ dyads, 690 UR dyads). Here, top and bottom rows reflect data acquired on different days of data collection while subjects viewed largely non-overlapping sets of movie clips. (B) Group-average ISC values used to create the difference maps in A, plotted in order of average ISC across all subject pairs, show that group differences are most pronounced in parcels with medium to high ISC (shading = SEM).

### Patterns of brain activity during movie-watching are heritable

After establishing that more genetically similar individuals share more similar movie-evoked BOLD time courses, we next sought to quantify the heritability of these brain responses. To do this, we leveraged a multidimensional estimator that has been used to assess the heritability of similar brain phenotypes (Anderson et al., 2021; Busch et al., 2023; Ge et al., 2016). Controlling for age, gender, and head motion, we found that movie-evoked BOLD time courses were heritable across almost all of cortex on both days of data collection (Fig. 2A, Day 1 mean *h*^2^ = .064 ± .034 (SD), Day 2 mean *h*^2^ = .068 ± .036, 99% of parcels significant on both days at FDR-corrected *P* < .05; dorsal/ventral views in Fig. S3A), and the spatial pattern of heritability across the cortex was very consistent across days of data collection (Spearman *ρ* = .96, *P*_BrainSMASH_ < .001). We note that we observed nearly identical results when we repeated this analysis without global signal regression (GSR; Fig. S4). Although heritability studies of one-dimensional traits (e.g., height) tend to require larger samples than the one used here, the multidimensional nature of our analysis affords us considerable power to detect small effects even with our relatively modest sample size (Anderson et al., 2021; Ge et al., 2016). To illustrate this point, we repeated our BOLD time course heritability analysis after excluding up to 90% of families in the present dataset and found that even after excluding half of our subjects, the average difference in *h*^2^ magnitude across parcels and between each subsample and the results reported above was less than .01, and the average spatial correlation (Spearman *ρ*) between subsample and full sample heritability values was greater than .9 (Fig. S5).

**Figure 2.**
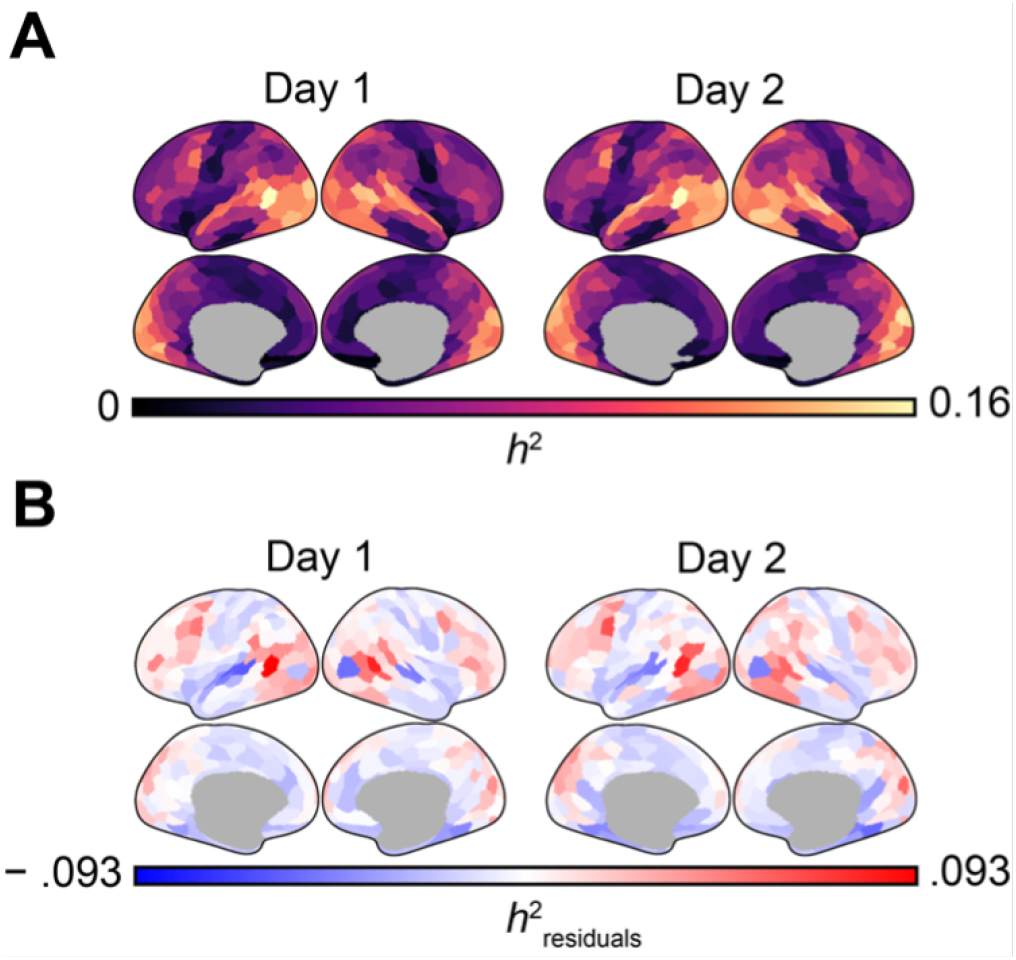
BOLD time courses are heritable across the cortex. (A) Cortical surfaces show heritability of BOLD time courses parcellated using the Schaefer 400 atlas, controlling for age, gender, and head motion (mean *h*^2^ Day 1 / Day 2 = .064 ± .034/ .068 ± .036). (B) Residuals after regressing parcel-level ISC from parcel-level heritability show that BOLD time courses in auditory cortices are less heritable than would be expected based on ISC, whereas the opposite is true for lateral prefrontal and temporo-parieto-occipital junction parcels.

Unsurprisingly, the spatial pattern of BOLD time course heritability was closely related to the spatial pattern of ISC (Day 1: Spearman *ρ* = .88, *P*_BrainSMASH_ < .001, Day 2: Spearman *ρ* = .86, *P*_BrainSMASH_ < .001), reflecting the simple fact that the heritability of movie-evoked BOLD time courses will be lower in parcels with less movie-driven activity to begin with. To characterize the heritability of BOLD time courses relative to the amount of movie-driven activity in each parcel, we regressed parcel-level ISC values (averaged across subject pairs) from heritability values and plotted the residuals in Fig. 2B (dorsal/ventral views in Fig. S3B). Here, negative values in the residual map indicate parcels where heritability is lower than expected based on ISC, while positive values indicate higher-than-expected heritability. Regarding alternative approaches to controlling for ISC, although the heritability model introduced by Ge et al. (2016) allows for the inclusion of covariates defined at the subject level (e.g., age), it does not allow for covariates that are defined at the dyad level (e.g., pairwise ISC). We observed that BOLD time courses were disproportionately more heritable in more associative lateral prefrontal and temporo-parieto-occipital junction parcels, while responses in lower-level auditory areas were less heritable than would be expected given their ISC. This indicates that although these more associative parcels do not encode more stimulus-specific information than unimodal sensory parcels, what information they do encode and/or how they encode it is under increased genetic control compared to auditory parcels.

After establishing that movie-evoked BOLD time courses are heritable, we next sought to determine the extent to which this heritability reflects genetic control over high-vs. low-level sensory processing. To do this, we leveraged the fact that low-level features of movie stimuli (e.g., visual motion and speech) tend to oscillate on the order of seconds (or faster), whereas higher-level aspects of the stimulus (e.g., social content and narrative structures) are encoded at lower frequencies (Baldassano et al., 2017; Honey et al., 2012; Kauppi et al., 2010). Similar to previous work on frequency-specific ISC (Kauppi et al., 2010), we filtered our data into five non-overlapping frequency bands, each containing an equal proportion of the total spectral power, and generated overall and residualized (with respect to ISC) heritability maps for each band (Fig. 3A and D). We observed that cortex-wide BOLD time course heritability increased monotonically with the period of the frequency band, such that heritability was over 50% higher in the slowest frequency band (0.004–0.02 Hz) compared to the unfiltered data (Fig. 3B and E). This suggests that genetic factors influence the neural processing of complex audio-visual features, and that this influence is greater than for lower-level sensory features. Interestingly, we also observed that both overall and residualized heritability were considerably lower in the one supra-BOLD frequency band (0.14–0.5 Hz, faster than the frequency of BOLD signals resulting from neuronal firing events; Josephs and Henson, 1999, Fig. 3A/D second column from the left) compared to the unfiltered data. This indicates that although there is synchronized high-frequency information in our data (possibly due to aliased cardiovascular and respiratory signals; Pérez et al., 2021), this information is largely not heritable and further supports a BOLD etiology for the heritability results shown above.

**Figure 3.**
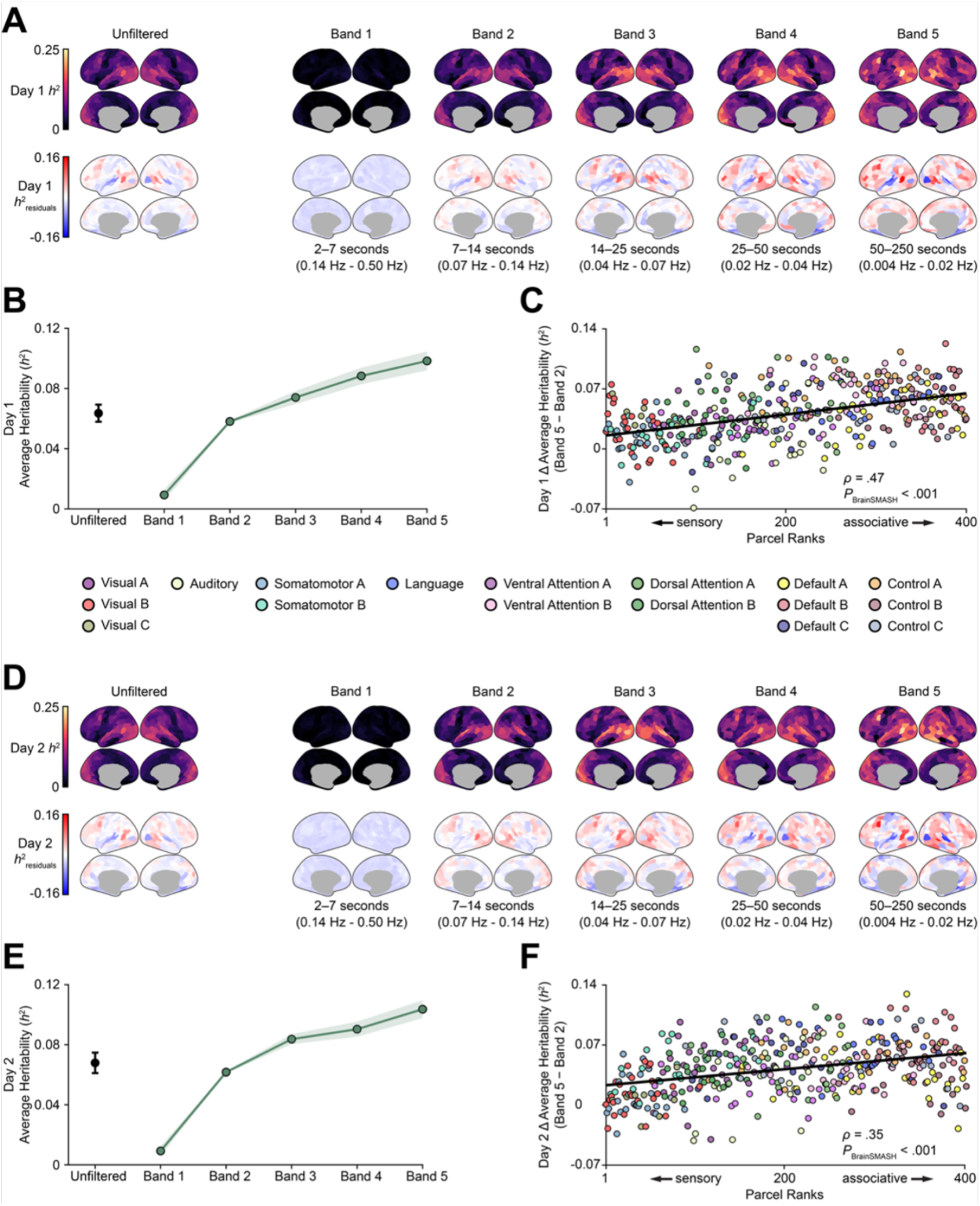
BOLD time course heritability is greater in slower frequency bands, especially for more associative parcels. (A) Purple/yellow cortical surfaces (upper row) show unfiltered BOLD time course heritability (upper left is identical to Fig. 2A) as well as the heritability of BOLD time courses filtered with five frequency bands, with greater heritability in slower bands for Day 1 data. Red/blue cortical surfaces show BOLD time course heritability residuals after regressing out parcel- and frequency-level differences in ISC (lower left is identical to Fig. 2B), with greater residuals in slower frequencies and more associative parcels. (B) Scatter plot shows heritability averaged across the cortex for each frequency band (i.e., the averages of the upper row of surfaces in A; shading = jackknife SEM). (C) Scatter plot shows the difference in heritability between the slowest and fastest BOLD-sensitive frequency bands for each of the Schaefer 400 parcels plotted against parcel ranks from the Sydnor et al. sensorimotor-association hierarchy (higher = more associative). Least squares lines were added to highlight the positive relationships between average *h*^2^ and parcel ranks but note that these relationships were formally tested with Spearman correlations. (D–F) Same as A–C for Day 2 data.

Previous studies have shown that during movie-watching, more associative regions process abstract information at longer timescales that range from tens of seconds to minutes, whereas sensory areas encode lower-level features at higher frequencies (Baldassano et al., 2017; Hasson et al., 2008; Honey et al., 2012). As such, we hypothesized that the higher heritability we observed in slower frequency bands was driven by increased heritability in associative (vs. sensory) parcels. To test this hypothesis, we correlated parcel-level differences in heritability between the slowest and fastest BOLD-sensitive frequency bands with sensorimotor-association hierarchy rankings from Sydnor et al. (2023; higher ranking = more associative) and found that heritability increases from the fastest to slowest frequency band were indeed larger for more associative parcels (Day 1: Spearman *ρ* = .47, *P*_BrainSMASH_ < .001, Day 2: Spearman *ρ* = .35, *P*_BrainSMASH_ < .001; Fig. 3C and F). Because removing rest and onset blocks from each clip and concatenating the two movie-watching runs from each day introduced temporal discontinuities that could impact our filtering results, we reran our analyses using the original, uncensored time courses and observed similar results (Fig. S6). We chose to initially analyze BOLD time courses parcellated using the Schaefer 400 atlas because parcellation reduces multiple comparisons, noise, and computational burden. However, we repeated our heritability analyses using data parcellated with 9 other resolutions of the Schaefer atlas and found that heritability reliably increased with average parcel size (Fig. S7). Moreover, the interpretation of parcellated BOLD time course heritability is complicated by the fact that macroscale areal boundaries are known to be heritable (Xu et al., 2016). As such, we use vertex-level data in our subsequent BOLD time course analyses.

### Heritable movie-evoked BOLD time courses reflect heritable cortical topographies

Our analyses of data aligned using standard anatomical methods (i.e., MSM) have demonstrated that patterns of movie-evoked brain activity and connectivity are heritable. Importantly, these patterns reflect two distinct and fundamental aspects of brain function: *how* stimuli are processed and *where* stimuli are processed. For example, when analyzing brain responses in a dorsal brain region (as shown in Fig. 4A), two twins (left and center) may appear to process stimulus information more similarly than an unrelated individual (right), based on having higher ISC values for that region. However, this apparent disparity arises purely from spatial differences in where the same information is processed: the unrelated individual in fact exhibits the same functional responses as the twins (i.e., the green and purple time courses), just in different cortical locations. These individual-specific maps of how shared brain functions are spatially distributed are known as cortical topographies (Haxby et al., 2011, 2020), and recent work has shown that cortical topographies defined at rest are influenced by genetic factors (Anderson et al., 2021; Burger et al., 2022; Busch et al., 2023). Therefore, we hypothesized that part of the heritability observed in our previous analyses might reflect genetic control over cortical topography (or “where” information is processed), in addition to genetic influences on information processing itself.

**Figure 4.**
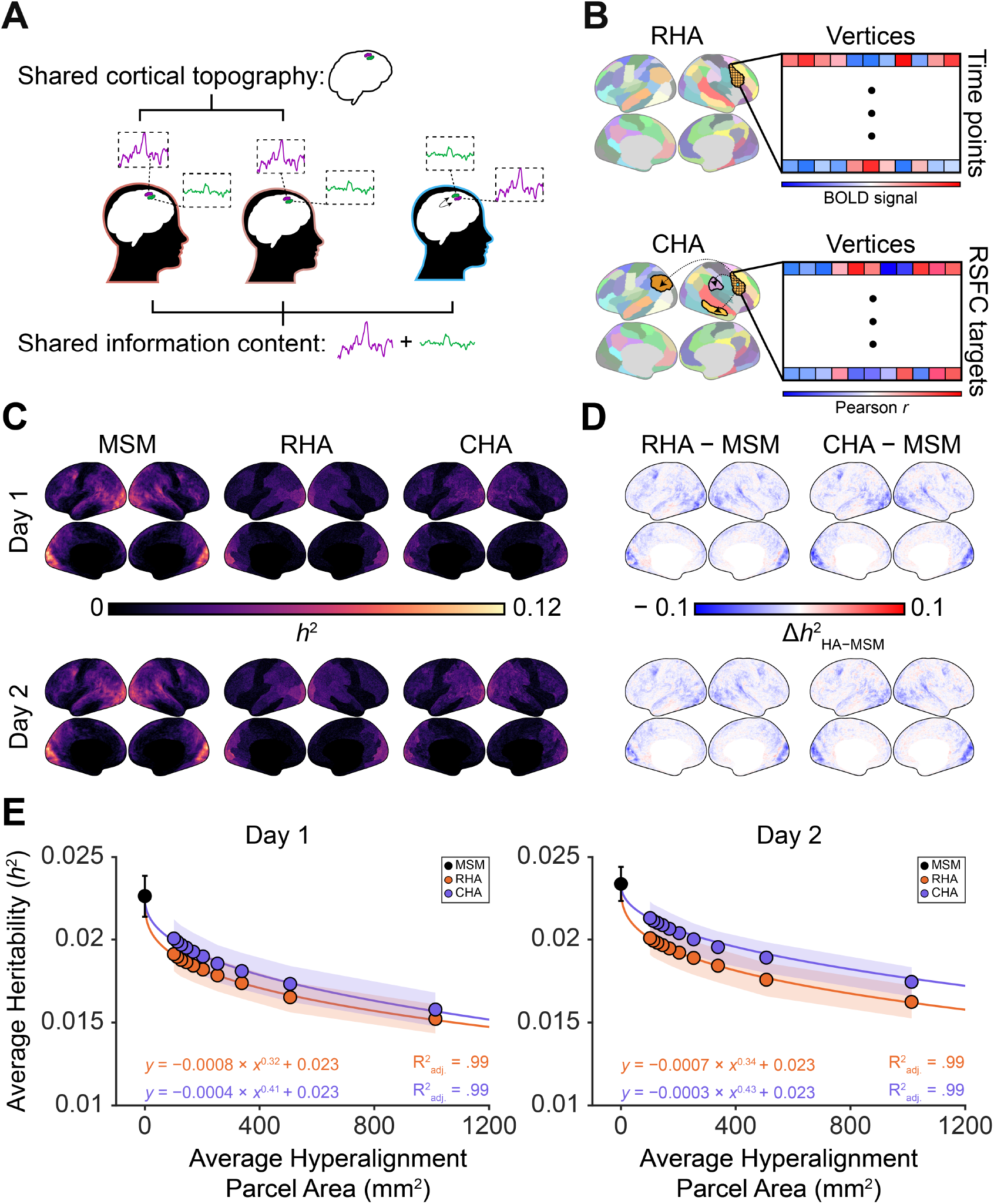
Hyperalignment reduces BOLD time course heritability. (A) Cartoon illustrates the difference between shared cortical topographies and shared (topography-independent) information content. (B) Diagrams illustrate the inputs to response and connectivity hyperalignment (RHA and CHA, respectively) using the Schaefer 100 atlas. RHA topographies were learned using BOLD time course data from the other day’s movie-watching scans, while CHA topographies were learned from vertex-level FC profiles (i.e., correlations between one vertex’s BOLD time course and the average time course from each of the 99 other parcels) calculated from the other day’s resting state scans. (C) Vertex-level BOLD time course heritability is highest for data aligned via MSM (multimodal surface matching) and lower for data hyperaligned within 100 Schaefer atlas parcels using both response hyperalignment (RHA) and connectivity hyperalignment (CHA). (D) Differences between the MSM-only and hyperaligned heritability maps shown in (C) are distributed across the cortex but are most apparent in visual areas. (E) BOLD time course heritability decreases as a function of hyperalignment parcel size according to a power law (purple and orange lines); each dot corresponds to average cortex-wide heritability for data hyperaligned using one of the 10 Schaefer atlas resolutions (shading = jackknife SEM).

One effective method used to separate the topography-dependent and topography-independent aspects of cortical information processing is known as hyperalignment. Hyperalignment aligns individual brains into a common high-dimensional functional space based on shared functional responses during the same task or stimulus (Haxby et al., 2011). By aligning fMRI data across subjects into a topography-independent functional space, hyperalignment yields datasets that allow for a direct comparison of how information is processed across individuals, independent of individual differences in where that information is processed.

In this study, we used hyperalignment to quantify the extent to which brain response heritability reflects genetic control over *how* vs. *where* information is processed. To hyperalign each subject’s movie-watching data to a common functional space for a given day of data collection, we used idiosyncratic transformation matrices that were learned from either the other day’s movie activity time courses (response hyperalignment, or RHA) or from rest FC profiles calculated from the other day’s scans (connectivity hyperalignment, or CHA). Although RHA and CHA align fMRI data with similar fidelity (Guntupalli et al., 2018; Haxby et al., 2020), using both methods allows us to evaluate whether heritable functional topographies reflect the brain’s intrinsic functional architecture or movie watching-specific response functions. We performed both RHA and CHA in a piecewise fashion (Bazeille et al., 2021), aligning vertex-level data within individual Schaefer parcels—in other words, a vertex in one Schaefer parcel would be aligned with other vertices in that parcel and never with vertices from other parcels.

Hyperalignment aligns vertices with similar functional responses across subjects, inherently increasing ISC across subject pairs (Fig. S8). However, to the extent that cortical topographies are under genetic control, twins’ brains are intrinsically more aligned than those of unrelated individuals. Therefore, we predicted that hyperalignment would decrease observed heritability across the cortex by eliminating the topography-dependent component of heritability and increasing response similarity more in unrelated dyads than in twin pairs.

Starting with the coarsest Schaefer atlas resolution (100 parcels, mean parcel area = 1,013 mm^2^), we found that RHA and CHA significantly decreased BOLD time course heritability to similar degrees across the cortex. Compared to MSM-aligned data (Fig. 4C, left column), hyperalignment reduced BOLD time course heritability across the cortex by 33% on Day 1 (95% CI = [25–40%]) and 31% on Day 2 [22–39%] for RHA (Fig. 4C, middle column), and by 30% [21–39%] and 25% [18–33%] for CHA (Fig. 4C, right column; dorsal/ventral views in Fig. S9). These decreases were most apparent in visual cortex, but were also prominent in associative areas like the right temporoparietal junction and bilateral area 55b (Fig. 4D), and the spatial pattern of this effect was consistent across days (RHA: Spearman *ρ* = .66, *P*_BrainSMASH_ < .001, CHA: Spearman *ρ* = .69, *P*_BrainSMASH_ < .001) and hyper-alignment methods (Day 1: Spearman *ρ* = .76, *P*_BrainSMASH_ < .001, Day 2: Spearman *ρ* = .74, *P*_BrainSMASH_ < .001).

To quantify the spatial scale at which cortical topography influences BOLD time course heritability, we then repeated RHA and CHA using the 9 other Schaefer atlas resolutions (200 to 1000 parcels). Because hyperalignment can eliminate more heritable differences in cortical topography when it is performed in larger parcels, we predicted that hyperalignment across larger parcels would decrease BOLD time course heritability to a greater extent. As expected, the magnitude of these reductions decreased as hyperalignment was performed across smaller areas (Fig. 4E). To quantify this relationship, we fit power law models of the form *y* = *a*⋅*x*^*b*^ +*c* to the 11 hyperalignment resolutions (corresponding to the average parcel areas for the 10 Schaefer atlases as well as 0 for no hyperalignment) and their corresponding average *h*^2^ values. We found that these power law models accurately characterized how heritability scaled with hyperalignment resolution for both RHA and CHA on Day 1 (RHA: *y* = −0.0008 ⋅ *x*^0.32^ + 0.023, R^2^_adj._ = .99, CHA: *y* = −0.0004 ⋅ *x*^0.41^ + 0.023, R^2^_adj._ = .99) and Day 2 (RHA: *y* = −0.0007 ⋅ *x*^0.34^ + 0.023, R^2^_adj._ = .99, CHA: *y* = −0.0003 ⋅ *x*^0.43^ + 0.023, R^2^_adj._ = .99), whereas linear, quadratic, and logarithmic models performed worse (Fig. S10).

### Heritability of BOLD time courses is related to neural timescales

In the previous section, we found that individual differences in a stable aspect of brain function (cortical topography) accounted for 30–40% of the heritability of movie-evoked brain responses. Importantly, cortical topography is a largely spatial trait. Although it captures significant inter-individual variability in how brain function varies over space, it is not directly related to how brain responses evolve over time. As such, we reasoned that the heritability of high-dimensional brain responses might also be grounded in the heritability of stable, temporal properties of brain function.

One such property is the neural timescale (NT), which is thought to index the duration of information storage in a given circuit or region. Across the cortex, more associative areas are known to have longer NTs (which is consistent with our frequency-dependent heritability results in Fig. 3), but substantial and behaviorally-relevant variability in NTs also exists across individuals, such that individuals with longer NTs in a given region integrate sensory information across longer periods of time (Wengler et al., 2020). NTs are commonly operationalized as the area under the curve of the autocorrelation function (ACF) until the lag preceding the first negative ACF value (Wengler et al., 2020). Because stimulus-evoked time courses that are more autocorrelated will tend to be more correlated with each other (see Methods), we suspected that NTs could be an important determinant of ISC. To test this, we correlated the sum of each dyad’s NTs from one day’s movie-watching scan with their ISC from the other movie-watching scan and found that we could explain a considerable portion of the variability in pairwise ISC from NT alone (max/mean correlation Day 1: *ρ* = 0.56/0.10, Day 2: *ρ* = 0.65/0.11, 54% of vertices significant at FDR-corrected *P*_perm_ < .05 on both days, Fig. S11). Given this relationship between NT and ISC as well as the fact that a similar measure of BOLD autocorrelation was recently shown to be heritable (Christova et al., 2022), we hypothesized that some of the BOLD time course heritability not accounted for by topography is underpinned by heritability of NT.

Before testing this hypothesis directly, we first sought to establish that NT magnitude and topography themselves are heritable. After calculating NTs at each vertex from each day’s movie-watching data, we averaged these values to generate a single cortex-wide NT (i.e., global NT) for each subject and each day of data collection. We then examined pairwise differences in global NT across the three groups introduced earlier: MZ twins, DZ twins, and age- and gender-matched UR dyads. As expected, we found that differences in MZ twins’ global NTs were smaller than in unrelated individuals’ NTs on both days (Fig. 5A, Day 1: ΔNT_MZ_ = 0.14 ± .015 (SD), ΔNT_DZ_ = 0.18 ± .022, ΔNT_UR_ = 0.23 ± .007; Day 2: ΔNT_MZ_ = 0.15 ± .013, ΔNT_DZ_ = 0.24 ± .032, ΔNT_UR_ = 0.26 ± .007). We then used SOLAR to formally quantify the heritability of this trait, finding that global NT was heritable on both Day 1 (*h*^2^ = 0.38, *P* = .003) and Day 2 (*h*^2^ = 0.75, *P* = 3.7 × 10^−9^). We note that the discrepancy between *h*^2^ effect sizes on Day 1 and Day 2 reflects decreased power to resolve the heritability of one-dimensional traits, as discussed above. Finally, we used a multidimensional heritability analysis to establish the heritability of parcel-level NT to-pographies, finding that NT topographies (or the scale-invariant pattern of NTs across 400 cortical parcels) was heritable at *h*^2^ = 0.39 on both Day 1 and Day 2 (*P*_perm_ < .0001 on both days).

**Figure 5.**
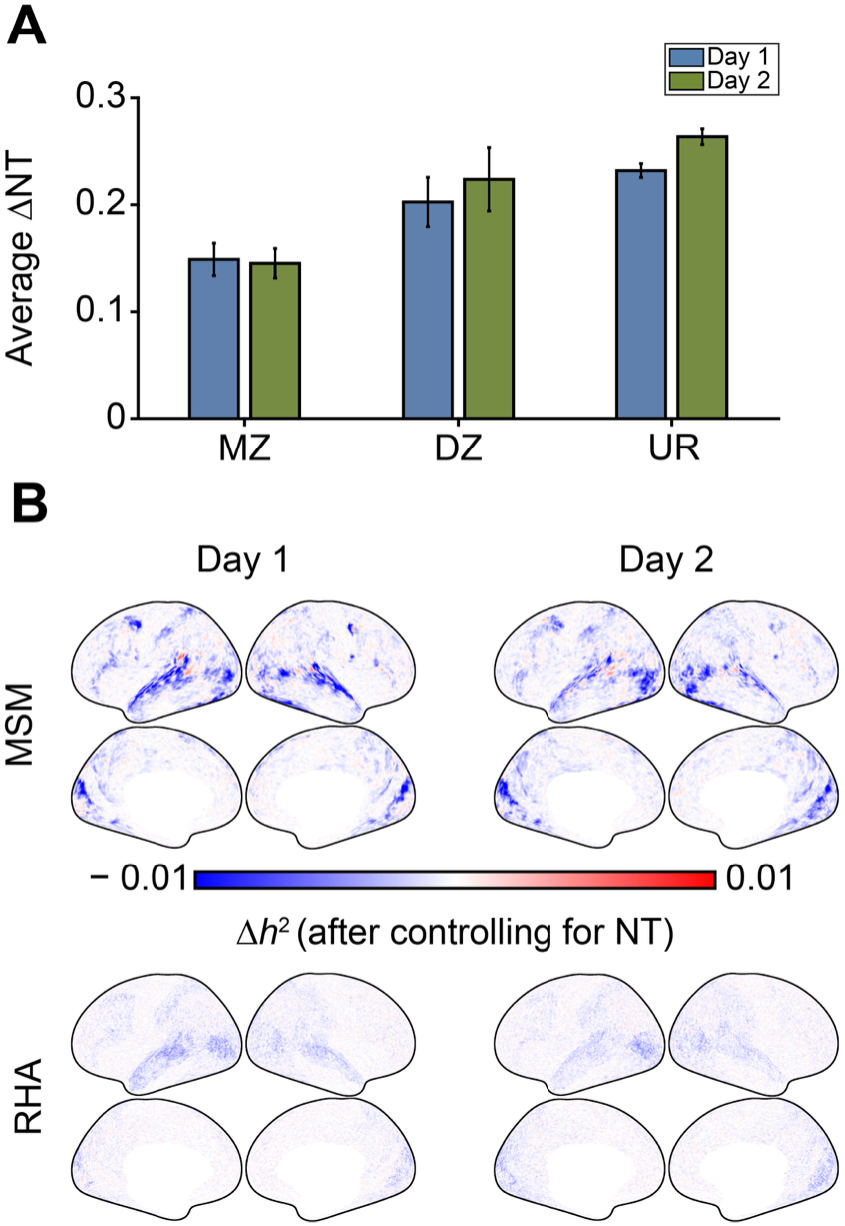
Controlling for neural timescale (NT) reduces heritability of BOLD time courses. (A) Bar plots show average pairwise differences in cortex-wide NT across MZ, DZ, and UR dyads on both days of data collection. (B) Cortical surfaces show decreases in BOLD time course heritability after NTs calculated from the other day of data collection were included as covariates in the multidimensional heritability analyses for MSM-aligned and RHA-aligned (using the Schaefer 100 parcellation) data, most prominently in mid-level auditory and visual regions. These maps are thresholded at Δ*h*^2^ = ±0.01 to aid comparisons of MSM- and RHA-aligned results. The maximum differences in *h*^2^ after controlling for NTs were −0.025 for MSM-aligned data and −0.007 for RHA-aligned data, respectively.

To quantify the extent to which BOLD time course heritability reflects genetic control over NT, we repeated the heritability analyses from earlier for each day of data collection, this time including vertex-level NTs calculated from the other day’s data as co-variates (in addition to age, gender, and head motion). We observed that controlling for NT significantly reduced BOLD time course heritability on both days in 5.5% of vertices (Fig. 5B upper row, FDR-corrected *P*_perm_ < .05 separately for each day; raw pre-hyperalignment voxel-level heritability maps available in Fig. S12), most prominently in speech and language areas (e.g., auditory/superior temporal cortices, area 55b) and motion-sensitive visual areas (e.g., medial temporal and medial superior temporal cortices). Although these reductions in heritability were more focal than those observed following hyperalignment, they reached similar strengths, with decreases of 20–30% observed throughout the superior temporal gyri on both days of data collection.

After establishing that cortical topographies and neural timescales both contribute to the heritability of high-dimensional brain responses, we next sought to determine if these contributions are independent of one another. To test this, we recalculated BOLD time course heritability following RHA using the Schaefer 100 parcellation (the most aggressive hyperalignment approach from the previous section), this time controlling for NT calculated from these RHA-aligned data. If cortical topography and neural timescale constitute separable processes through which genetics shapes brain responses, we would expect controlling for NT in the RHA- and MSM-aligned data to reduce brain response heritability to similar degrees. Instead, we observed a mixed result: although controlling for NT after hyperalignment further reduced BOLD time course heritability on both days in 1.5% of vertices (FDR-corrected *P*_perm_ < .05 separately for each day, Fig. 5B, lower row), the average magnitude of this decrease across the cortex was 40% smaller than for the MSM-aligned data. This suggests that cortical topography and neural timescale each account for some unique variance in brain response heritability, but their contributions are not entirely independent.

### Movie-evoked FC profiles are heritable and reflect heritable cortical topographies

Thus far, we have demonstrated that movie-evoked BOLD time courses are heritable, and that this heritability is related to genetic control over stable spatial and temporal aspects of brain function. However, sensory information is encoded and processed not just in the activities of single regions but also in the functional connectivity (FC) between multiple regions (Chen et al., 2014). To determine if this other kind of movie-evoked brain response is similarly heritable, we repeated our analyses using movie-watching FC (movie FC) profiles. Here, a given individual’s movie FC profiles were calculated for each pair of 17 Kong networks and each day of data collection as their set of movie FC values for all connections between parcels in that pair of networks.

Starting again with a dyadic similarity analysis, we observed that for all comparisons, the more genetically related the dyads, the more similar their movie FC patterns (Fig. 6A, MZ>UR / MZ>DZ / DZ>UR: 49% / 26% / 18% greater similarity across network combinations). Compared to group differences in ISC, movie FC profile similarities showed greater separation between groups (Fig. 6B, group average FC profile similarity shown in Fig. S13).

**Figure 6.**
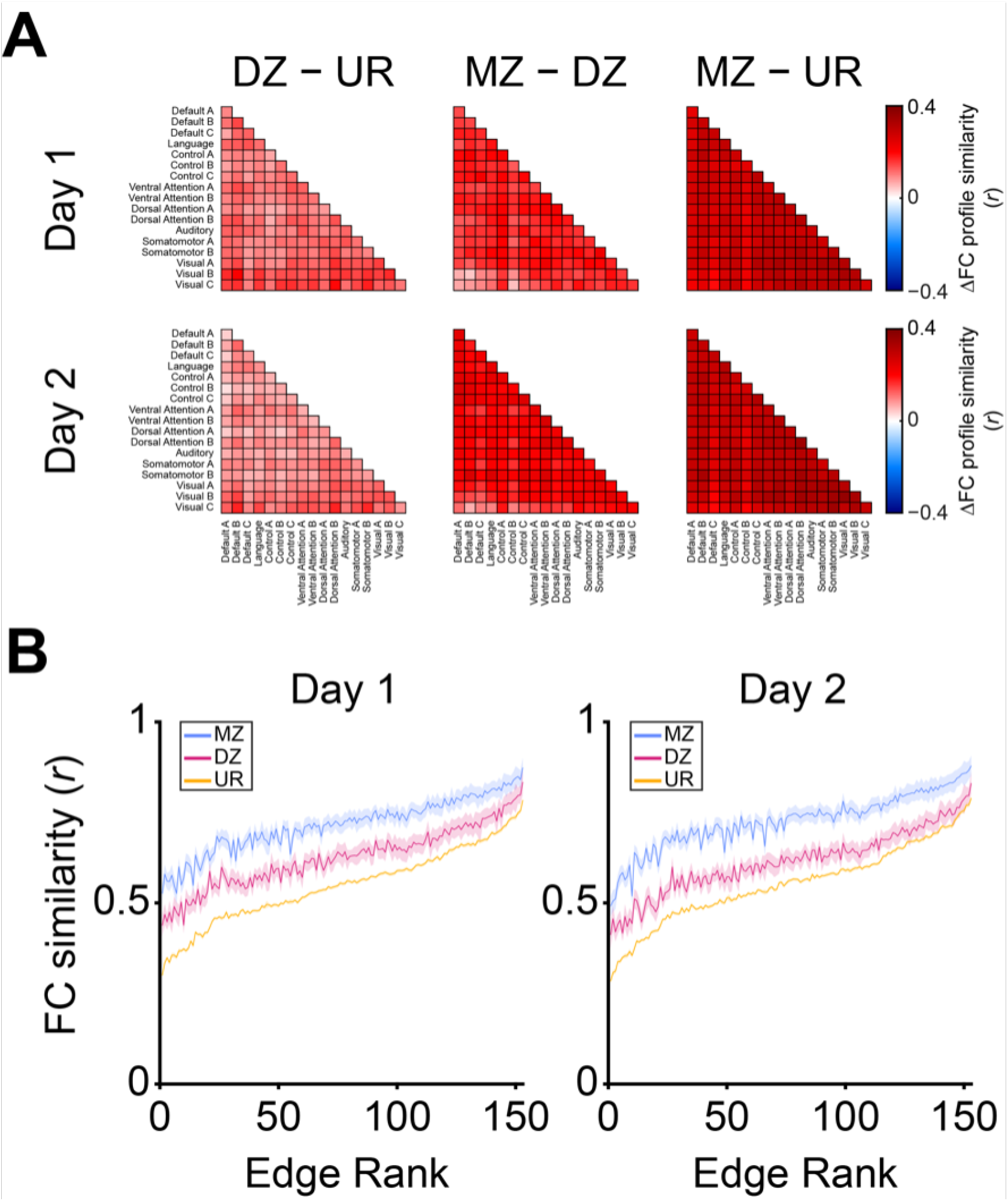
FC profile similarity scales with genetic relatedness across the cortex. (A) Group differences in average FC profile similarity show that FC profiles are more similar for dyads who are more genetically related (51 MZ dyads, 34 DZ dyads, 690 UR dyads). (B) Group-average FC profile similarity values used to create the difference maps in A, plotted in order of average FC profile similarity across all subject pairs (shading = SEM).

Given the greater between-group separation in FC profile similarity (vs. BOLD time course similarity) in Fig. 6B vs. Fig. 1B, we expected movie FC profiles to be more heritable than movie-evoked BOLD time courses. Applying the same multidimensional heritability analysis to movie FC profiles, we indeed found that FC profiles were around six times as heritable as BOLD time courses (Fig. 7, left column, Day 1 mean *h*^2^ = .36 ± .035, Day 2 mean *h*^2^ = .37 ± .038, all network combinations significant on both days at FDR-corrected *P* < .05), and the pattern of heritability values across network combinations was very consistent between days (Spearman *ρ* = .92, *P*_perm_ < .001). The six-fold higher heritability of multidimensional FC profiles compared to BOLD time courses likely results from each FC profile dimension representing a connectivity strength calculated over many time-points between two cortical areas, whereas each BOLD time course dimension reflects an activity magnitude at a single timepoint in one cortical area. Consequently, FC profile dimensions are less noisy (due to being defined across many timepoints) and capture more individual differences (due to being defined across multiple areas).

**Figure 7.**
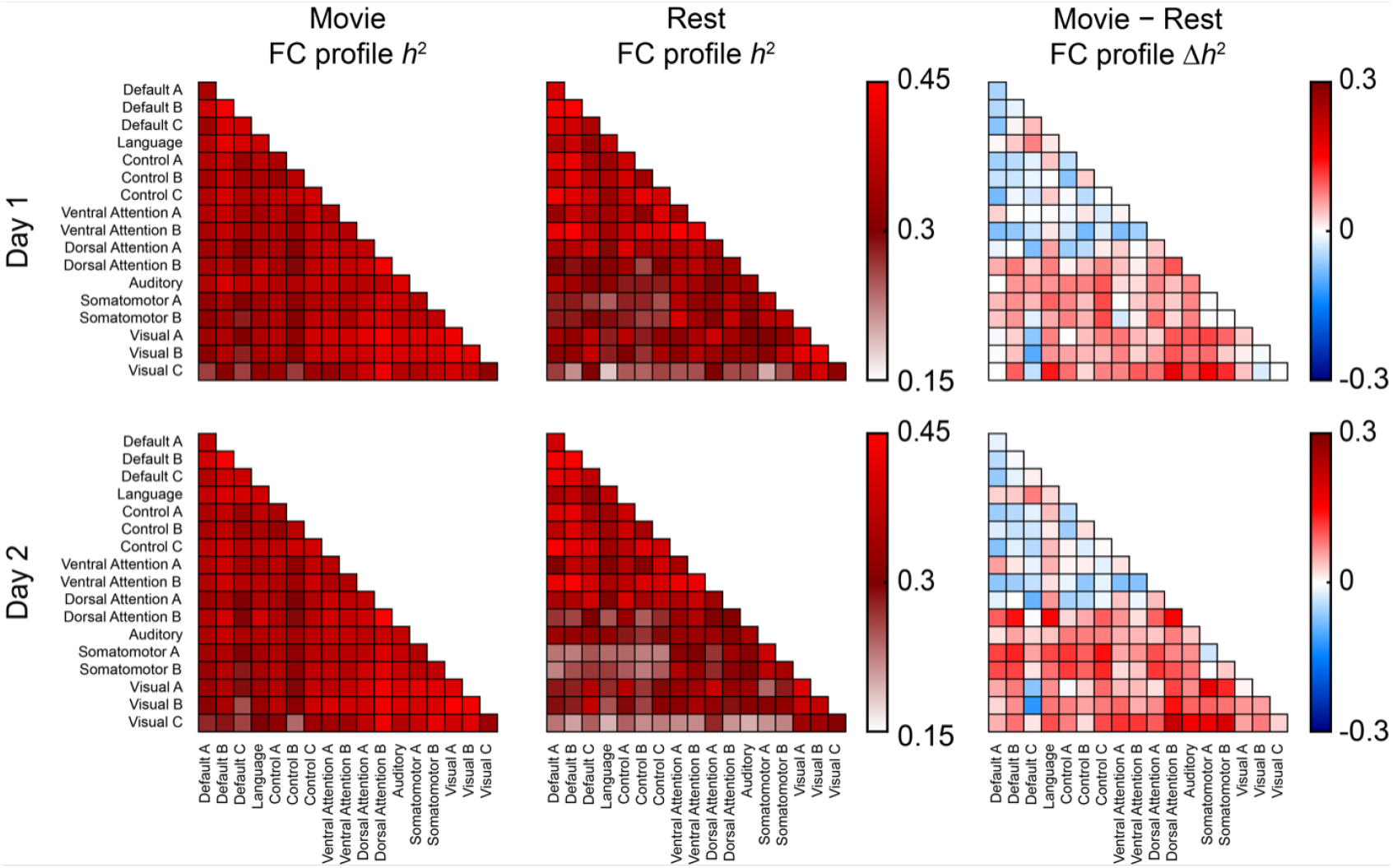
FC profiles are heritable across network combinations. Heatmaps show heritability of FC profiles for all unique within- and between-network combinations of the 17 Kong networks after controlling for age, gender, and head motion. FC profiles during movie-watching (left column) were more heritable than resting state FC profiles (middle column) for more sensory-oriented networks (red rows in the right column).

Although we are unaware of any previous studies that have investigated FC heritability during movie watching, the heritability of resting state FC (rest FC) measures has been well established (Anderson et al., 2021; Busch et al., 2023; Glahn et al., 2010; Miranda-Dominguez et al., 2018). Compared to rest FC, movie FC profiles serve as better identifiers of individuals (Vanderwal et al., 2017) and better predict individual differences in behavior (Finn & Bandettini, 2021). Given movie FC’s increased sensitivity to individual variability, we reasoned that movie FC profiles would be more heritable than rest FC profiles. Using resting state data collected from the same subjects and on the same days of data collection, we calculated rest FC profile heritability using the approach described above and found that heritability was indeed lower than for movie FC profiles (Fig. 7, right column), but this effect was limited to more sensory-oriented networks (Language, Auditory, Somatomotor A and B, Visual A, B, and C, and Dorsal Attention B networks significant at FDR-corrected *P*_perm_ < .05 on both days). Across these networks, movie (vs. rest) FC profiles were 20% more heritable on Day 1 (min. = 11%, max. = 30%) and 30% more heritable on Day 2 (min. = 19%, max. = 44%). Rest FC profiles were not significantly more heritable than movie FC profiles in any network. Because subjects tend to move more during resting state vs. movie-watching scans, and because this could explain the higher heritability of movie FC profiles, we repeated our analyses after censoring all frames with FD >0.2 mm and found the with- and without-censoring results to be nearly identical (data not shown).

Our connectivity analyses thus far have focused on FC profiles (i.e., correlations across FC values) instead of FC strengths (i.e., the average of FC values), as the low power afforded by our sample size precludes us from measuring the heritability of one-dimensional phenotypes with high precision (Benson et al., 2022; Busch et al., 2023). However, FC strengths are clinically and behaviorally relevant, and measuring how task conditions and alignment approaches affect FC strength heritability across the cortex requires substantially less power than resolving FC strength heritability values for individual network combinations. With this in mind, we used SOLAR (Almasy & Blangero, 1998) to estimate the heritability of FC strength in this and subsequent analyses and provide the corresponding figures in the Supplementary Materials (Fig. S14).

We next repeated our FC profile (and strength) analyses using the hyperaligned data. Similar to the effects of RHA and CHA on BOLD time course heritability, hyperalignment within the Schaefer 100 parcels significantly reduced FC profile heritability across network combinations by 39% on Day 1 (95% CI = [32–46%]) and 41% on Day 2 [34–48%] for RHA, and by 20% [13–28%] and 18%

[12–25%] for CHA (Fig. 8A). However, we did observe some consistent differences between effects of hyperalignment on FC profile vs. BOLD time course heritability. First, RHA decreased FC profile heritability to a significantly greater extent than did CHA, seen in the larger separation between purple and orange traces in Fig. 8B. Second, hyperalignment’s effects on FC profile (vs. BOLD time course) heritability were less variable across the different parcellation resolutions, seen in the relatively flat slope between orange/purple dots. This is reflected in the lower power law *b* coefficient values for FC profile (0.08–0.11, Fig. 8B) vs. BOLD time course heritability (0.32–0.43, Fig. 4E). We observed a similar area-independent pattern of results for our FC strength analyses, although here only RHA (and not CHA) significantly decreased FC strength heritability (Fig. S15). Once again, linear, quadratic, and logarithmic models failed to explain the relationships between hyperalignment resolution and FC profile (Fig. S16) and strength (Fig. S17) heritability compared to power law models.

**Figure 8.**
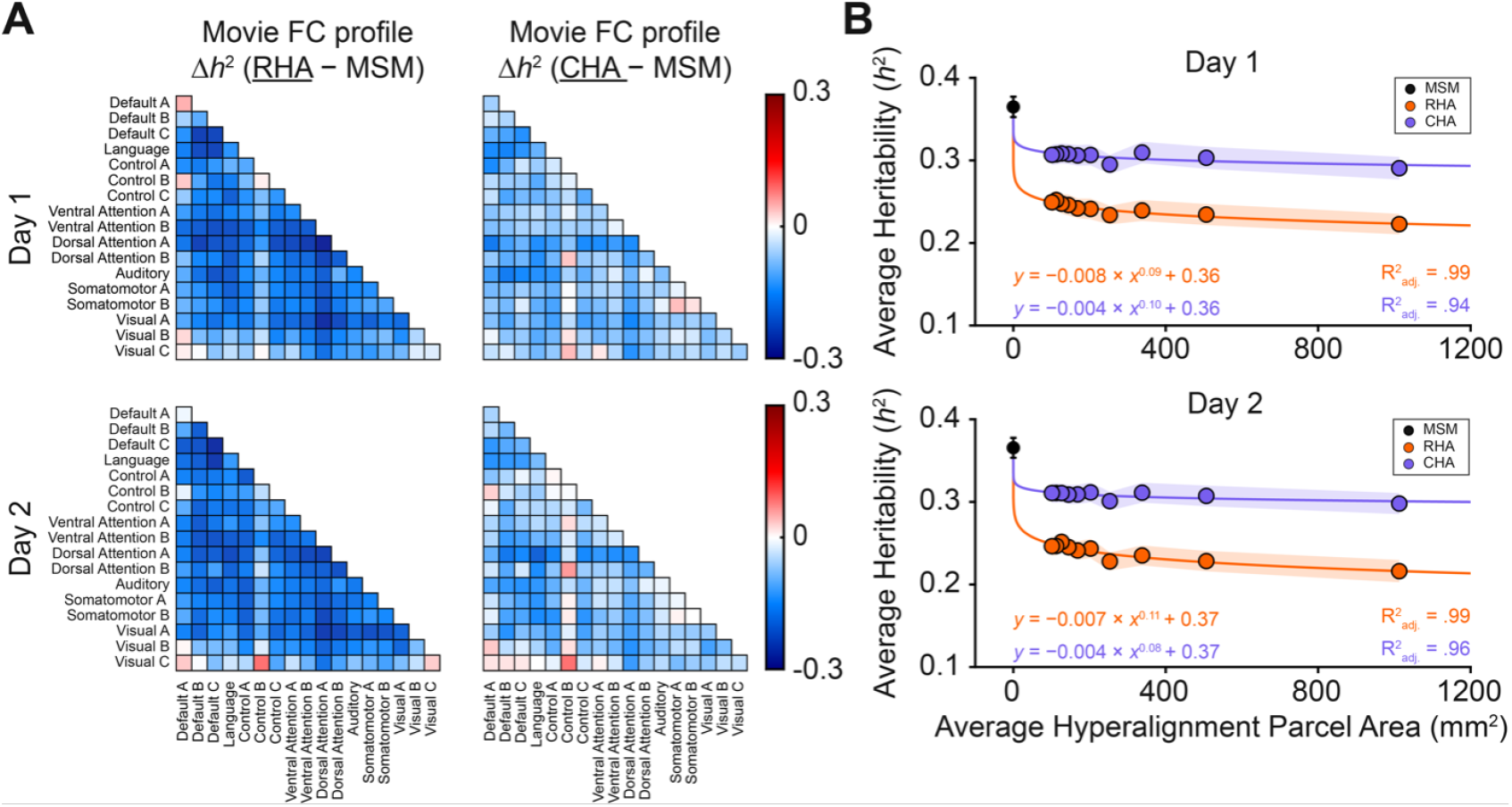
Hyperalignment reduces FC profile heritability. (A) Heatmaps show decreased FC profile heritability for most combinations of 17 Kong networks following RHA (left) and CHA (right) compared to the MSM-only baseline. (B) Scatter plots show that hyperalignment, especially with RHA, decreases FC profile heritability according to a power law function; each dot corresponds to average cortex-wide heritability for data hyperaligned using one of the 10 Schaefer atlas resolutions (or MSM-only alignment, shading = jackknife SEM).

## Discussion

In this study, we examined the heritability of movie-evoked BOLD activity and connectivity. First, we showed that BOLD time courses and FC profiles are heritable across the cortex, especially in and between the sensory and associative regions that are most reliably activated by the stimuli. Second, we showed that this heritability is underpinned by genetic control over fundamental spatial and temporal characteristics of brain function that reflect both *where* and *how* individuals process sensory information. More specifically, our findings demonstrate that genetics influences cortical topography as a power law function of cortical area, and that a key property of brain function—the neural timescale—is responsible for an additional portion of BOLD time course heritability, especially in auditory and speech-sensitive areas. Just as importantly, these results suggest a modest ceiling for how much of this stimulus-driven activity and connectivity is under genetic control, leaving the rest to non-genetic individual variation.

Studies using ISC to examine similarity of movie-evoked BOLD activity typically find highly conserved responses in auditory and visual areas. We found that more genetically related individuals exhibited greater ISC not only in these sensory areas, but also across most of cortex. This increased ISC could reflect more similar stimulus processing in a number of ways. For example, high-level attentional effects, such as twins attending to more similar aspects of the stimulus, could account for this increase (Ki et al., 2016; Song et al., 2021). Such an attentional effect would explain our finding that BOLD time courses in auditory cortex (vs. mid-level visual and oculomotor areas) were less (vs. more) heritable than would be expected based on their overall ISC (Fig. 2B), as eye movements gate incoming visual information and associated neural representations (Borovska & de Haas, 2024) but no analogous mechanism exists in the auditory system, and eye movement patterns during complex scene viewing are themselves moderately heritable (Kennedy et al., 2017). Alternatively, low-level stimulus processing effects, such as twins having more similar population tuning than non-twins, could also lead to greater ISC.

Our frequency-dependent heritability results offer some preliminary insights into the specific aspects of sensory processing that these shared activity patterns represent. Here, we observed that BOLD time course heritability was over 50% greater in the slowest frequency band compared to the unfiltered data, and that this effect was driven by increased low-frequency heritability in more associative parcels. Because these regions and frequency bands encode more abstract stimulus features, this result suggests that the neural processing of high-vs. low-level sensory information is under greater genetic control. Importantly, we note that interpretation of this result is limited by reverse inference, and future studies that directly modulate low- and high-level stimulus information will be necessary to more conclusively answer this question.

Our approach differs from previous studies of stimulus- or task-driven brain activity heritability in that our ISC-based analyses don’t require assumptions about the nature of neural responses that, if inaccurate, could decrease the sensitivity of heritability estimates. Furthermore, instead of collapsing brain activity measurements across trials or epochs, our analyses exploit the high-dimensional nature of BOLD time courses by considering the unique information present at each timepoint. This multi-dimensional aspect of our analyses allowed us to leverage the significant amount of data available per subject to detect reliable (spatial *ρ* > .9 for heritability maps across days) effects even at the level of individual vertices. Furthermore, although our BOLD time course heritability effect sizes were modest (*h*^2^ ≤ .25 for parcels, *h*^2^ ≤ .12 for individual vertices), we note that these are commensurate with other twin-based heritability estimates of sensory phenotypes measured with fMRI.

Compared to our activity-based analyses, our FC analyses were more in line with previous work. Indeed, the heritability of FC profiles has been investigated on multiple occasions, sometimes with the same multidimensional estimator of heritability used here (Busch et al., 2023; Elliott et al., 2019; Miranda-Dominguez et al., 2018). Still, several aspects of our study allowed us to reveal novel results and add new context to established findings. For example, by analyzing resting state and movie-watching data from the same subjects, we were able to show that FC profiles involving sensory-oriented networks were significantly more heritable during movie-watching than at rest. This finding is consistent with reports that movie FC profiles better identify individuals (Vanderwal et al., 2017) and predict variability in behavioral traits (Finn & Bandettini, 2021) than resting state FC profiles, and that including task data increases estimates of FC profile heritability (Elliott et al., 2019). Relatedly, recent work by Luppi et al. (2025) has shown that FC uniqueness decreases under greater levels of anesthesia, suggesting that resting-state scans, where participants may likewise drift toward reduced arousal states, could mute idiosyncratic (and thus heritable) variation that remains robust when attention is held by an engaging movie stimulus.

Although BOLD time courses and FC profiles often serve as the fundamental brain phenomena to be studied in fMRI experiments, they are complex entities that are underpinned by a variety of lower-level biological processes (Hillman, 2014). As such, the heritability of movie-evoked brain responses established here likely reflects genetic control over more basic aspects of brain function. We demonstrated how two of these aspects, cortical topography and neural timescale, contribute to the heritability of stimulus-driven BOLD activity.

First, we found that controlling for idiosyncratic cortical topographies via response and connectivity hyperalignment (RHA and CHA) decreased activity heritability across the cortex. This decrease was bigger when data were aligned across larger parcels, but the rate of this decrease slowed as a power law function of parcel area, illustrating that genetic control of cortical topography is greatest at the fine scale. In addition to decreasing BOLD time course heritability, we found that RHA and CHA also decreased FC profile heritability, echoing recent work showing that CHA decreases rest FC profile heritability in a developmental population (Busch et al., 2023). Compared to our activity-based analyses, we noticed a far weaker effect of hyperalignment resolution on FC profile heritability, likely because this analysis was performed at the parcel level and across spatially distributed brain networks (thereby reducing the impact of local functional alignment). Although hyperalignment served to reduce heritable individual variability across our analyses, the residual post-hyperalignment heritability might be more behaviorally relevant, as hyperalignment has been shown to increase associations between FC profiles and cognitive test scores (Feilong et al., 2021). With this in mind, future studies investigating genetic correlations between brain function and behavioral variables may benefit from hyperalignment, as it can factor out individual-specific cortical topography and thus yield more precise estimates of functional heritability.

Just as cortical topographies spatially constrain individual responses to incoming stimuli, neural timescales (NTs) are stable temporal features of brain function that shape high-dimensional activity and connectivity patterns (Shinn et al., 2023; Wengler et al., 2020). More specifically, longer NTs are thought to reflect greater recurrent excitation at the micro-circuit level and yield more stable integration of sensory information (Cavanagh et al., 2020; Watanabe et al., 2019). Here, we found that MZ twins had more similar NTs than unrelated dyads; the fact that NTs measured by fMRI track electrophysiological activity (Watanabe et al., 2019) suggests that this reflects similarities in how movie stimuli were neurally encoded. We next showed that this genetic similarity in NT magnitude contributes to genetic similarity in movie-evoked BOLD time courses, such that controlling for vertex-level NT accounted for up to ∼30% of BOLD time course heritability, an effect that was strongest in speech-related areas like the superior temporal gyri. Importantly, controlling for NT had a weaker effect on BOLD time course heritability after hyperalignment. This is evidence that the topographic distribution of NTs, over and above their magnitude, is under genetic control. Beyond genetics, our finding that subjects with longer NTs had more correlated movie-evoked BOLD time courses suggests that decreased ISC in patients with schizophrenia (Patel et al., 2021; Tu et al., 2019), autism (Salmi et al., 2013), and depression (Gruskin et al., 2020) may be underpinned by shorter NTs in these same populations (Watanabe et al., 2019; Wengler et al., 2020; Zheng et al., 2024).

Our work should be considered in light of an important demographic limitation. Almost 90% of subjects in the present sample identified as White, and all subjects were between the ages of 22 and 36. As heritability estimates are known to differ across populations and age groups (Schmitt et al., 2014; Zhang et al., 2023), the generalizability of our findings is limited by the demographic characteristics of the HCP Young Adult sample used here (Ricard et al., 2023). In spite of this limitation, this work constitutes an important first link between the growing fields of neuroimaging genetics and “naturalistic” neuroscience. By considering BOLD time courses and FC profiles alongside cortical topographies and neural timescales derived from independent data, we reveal a multi-layered genetic influence that extends from basic features of brain function to complex, individual-specific sensory processing patterns. This comprehensive approach paves the way for future research to dissect the biological mechanisms that link genetics with sensory processing in both typical and atypical populations. Finally, we note that less than half of the inter-individual variability we observed in movie-evoked BOLD time courses and FC profiles was heritable, leaving the majority of this variability to be explained by gene-environment interactions as well as non-genetic factors such as life experiences and current behavioral state. As such, additional work will be necessary to characterize these non-genetic factors.

## Acknowledgments

We thank Avram Holmes and Erica Busch for helpful conversations regarding this project. Author D.C.G. was supported by an NIH MSTP training grant (T32GM007367). Data were provided by the Human Connectome Project, WU-Minn Consortium (Principal Investigators: David Van Essen and Kamil Ugurbil, 1U54MH091657) funded by the 16 NIH Institutes and Centers that support the NIH Blueprint for Neuroscience Research; and by the McDonnell Center for Systems Neuroscience at Washington University. Author G.H.P. receives income and equity from Pfizer, Inc. through family. Authors D.C.G. and D.J.V. have no competing interests to declare. This preprint was created using the LaPreprint template (https://github.com/roaldarbol/lapreprint) by Mikkel Roald-Arbøl .

## Citation diversity statement

Recent work in several fields of science has identified a bias in citation practices such that papers from women and other minority scholars are under-cited relative to the number of such papers in the field (Bertolero et al., 2020; Caplar et al., 2017; Chatterjee & Werner, 2021; Dion et al., 2018; Dworkin et al., 2020; Fulvio et al., 2021; Maliniak et al., 2013; Mitchell et al., 2013; Wang et al., 2021). Here we sought to proactively consider choosing references that reflect the diversity of the field in thought, form of contribution, gender, race, ethnicity, and other factors. First, we obtained the predicted gender of the first and last author of each reference by using databases that store the probability of a first name being carried by a woman (Dworkin et al., 2020; Zhou et al., 2020). By this measure and excluding self-citations to the first and last authors of our current paper, our references contain 7.46% woman(first)/woman(last), 5.97% man/woman, 28.36% woman/man, and 58.21% man/man. This method is limited in that a) names, pronouns, and social media profiles used to construct the databases may not, in every case, be indicative of gender identity and b) it cannot account for intersex, non-binary, or transgender people. Second, we obtained predicted racial/ethnic category of the first and last author of each reference by databases that store the probability of a first and last name being carried by an author of color (Ambekar et al., 2009; Chintalapati et al., 2023). By this measure (and excluding self-citations), our references contain 13.38% author of color (first)/author of color(last), 15.92% white author/author of color, 22.39% author of color/white author, and 48.32% white author/white author. This method is limited in that a) names and Florida Voter Data to make the predictions may not be indicative of racial/ethnic identity, and b) it cannot account for Indigenous and mixed-race authors, or those who may face differential biases due to the ambiguous racialization or ethnicization of their names. We look forward to future work that could help us to better understand how to support equitable practices in science.

## Author contributions

David C. Gruskin: Conceptualization, Methodology, Formal Analysis, Visualization, Writing–Original

Draft, Writing–Review & Editing

Daniel J. Vieira: Code review, Writing–Review & Editing

Jessica K. Lee: Supervision, Writing–Review & Editing

Gaurav H. Patel: Conceptualization, Writing–Review & Editing, Supervision

