## Supplemental Materials for "Heritability of movie-evoked brain activity and connectivity"

### Supplementary Materials

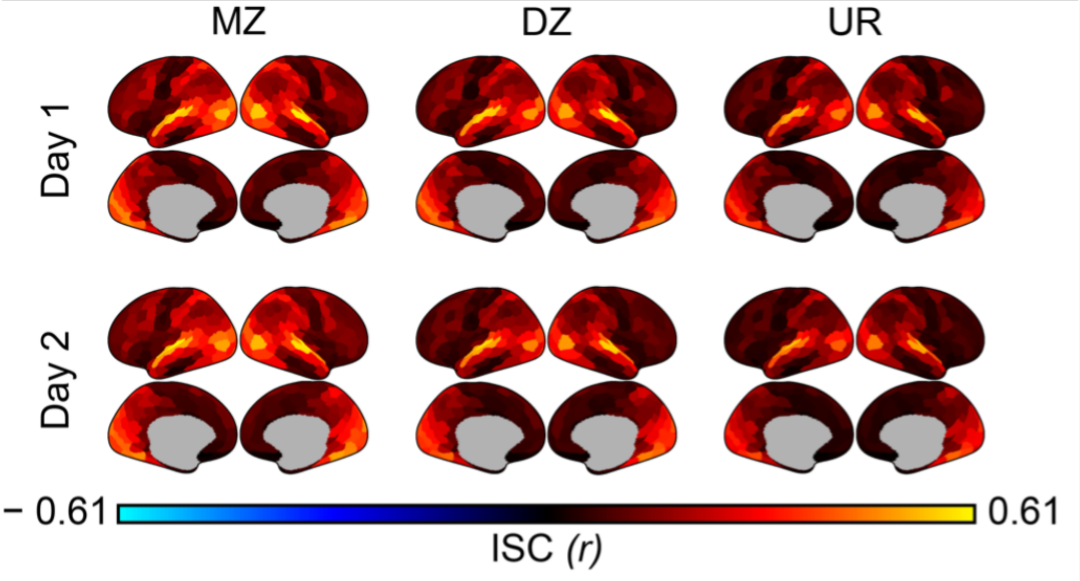


**Figure S1.** BOLD time course similarity by group. Cortical surfaces show the group-level ISC maps used to create the group difference maps in Fig. 1A

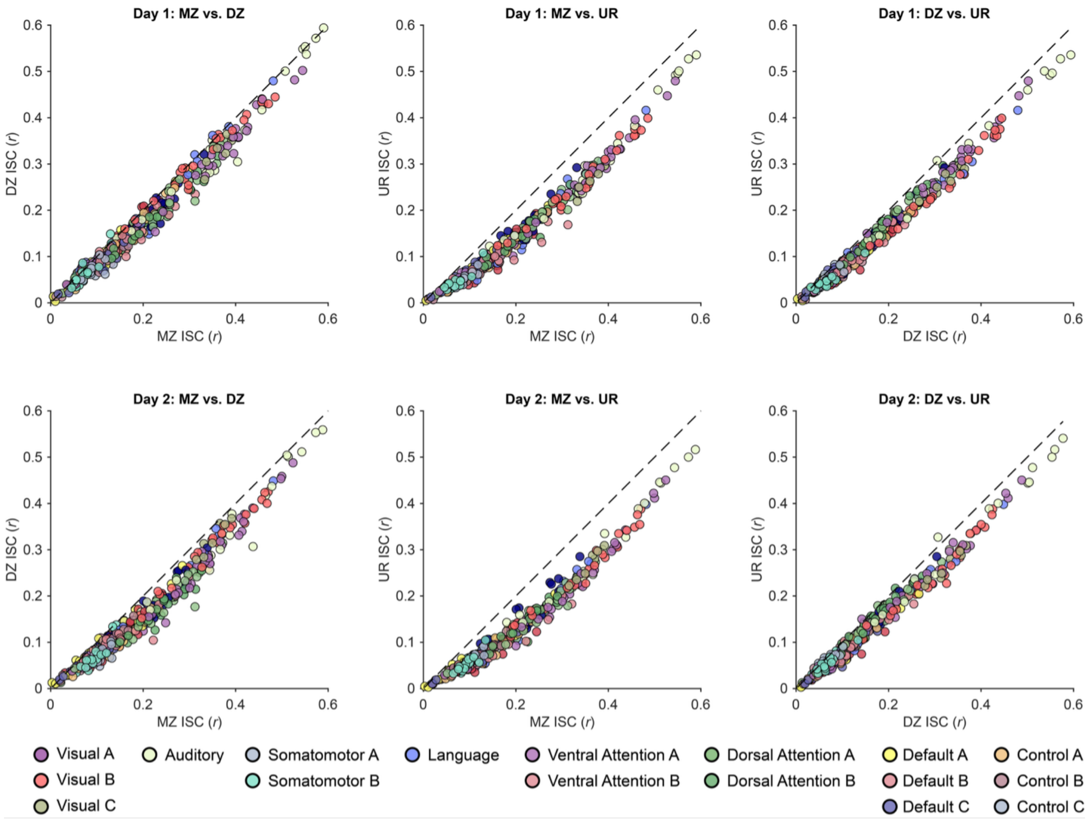


**Figure S2.** BOLD time course similarity scales with genetic relatedness across the cortex. Scatterplots show the same data as in Fig. 1B for each group comparison (each dot is one of 400 Schaefer parcels), highlighting greater genetic similarity in parcels with medium-to-high ISC.

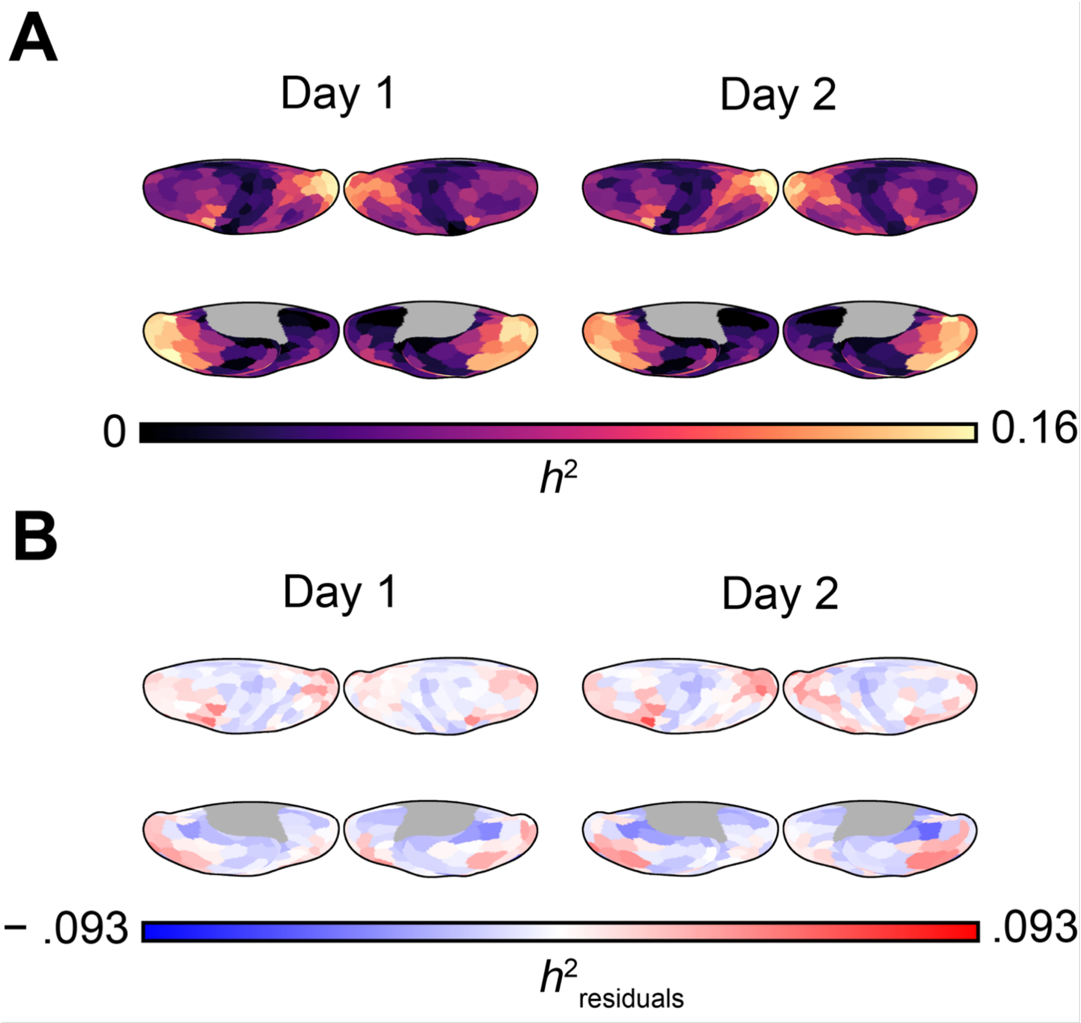


**Figure S3.** BOLD time courses are heritable across the cortex. Dorsal and ventral views of the same surfaces shown in Fig. 2

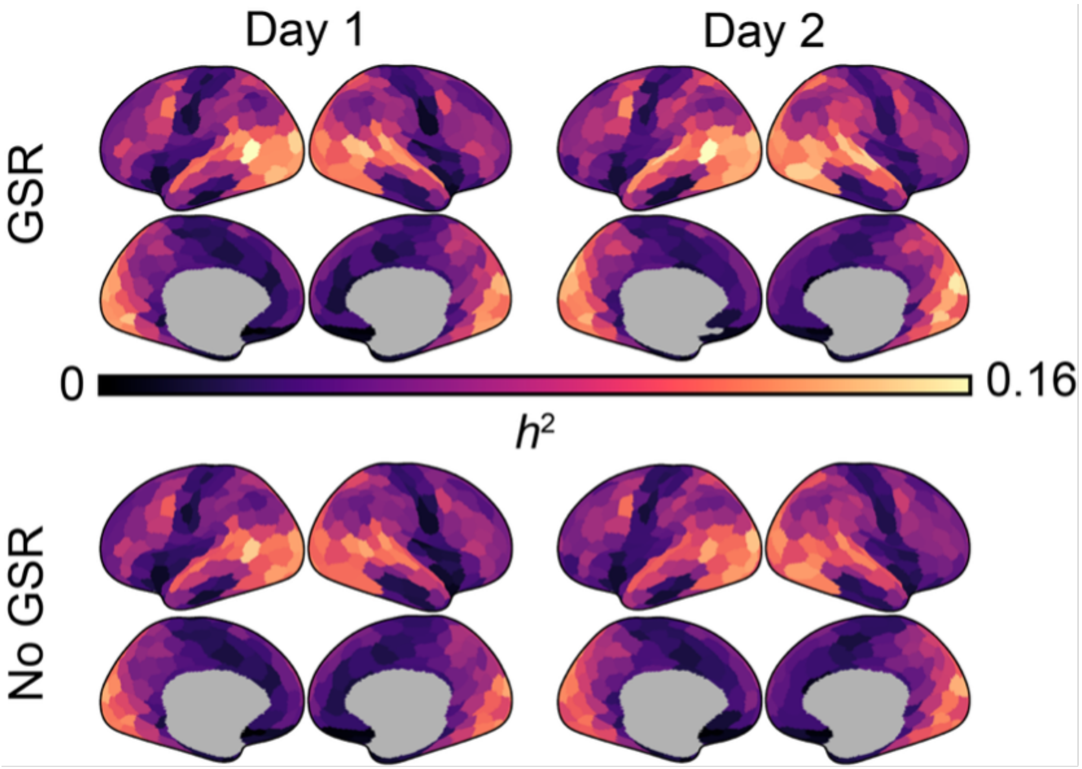


**Figure S4.** BOLD time course heritability is largely unaffected by GSR. Cortical surface show that GSR mildly increased BOLD time course heritability (average Day 1 $h^2$ with/without GSR = .064/.060; Day 2: .068/.061) and had almost no effect on its spatial pattern (With GSR/without GSR Spearman $\rho$ = .99, $P_{\text{BrainSMASH}}$ < .001 on both Day 1 and Day 2).

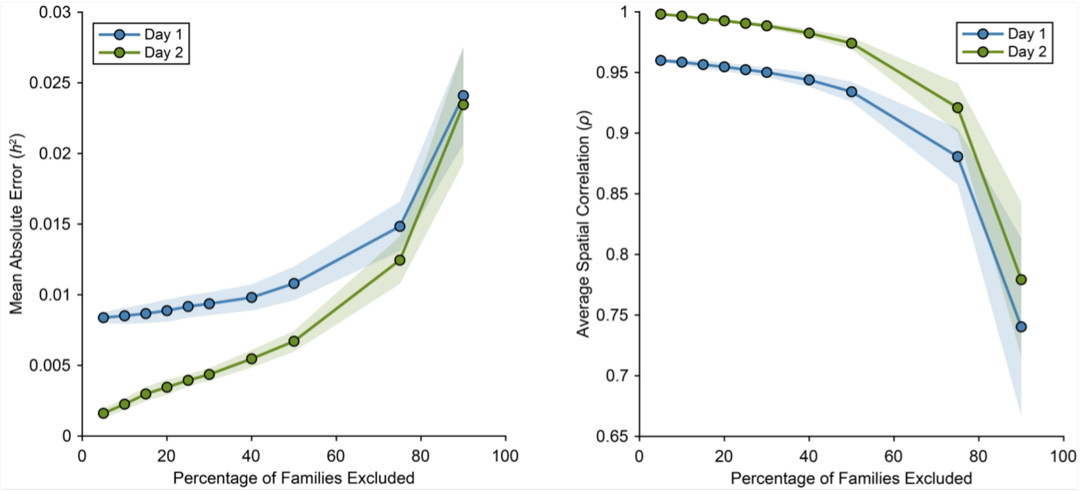


**Figure S5.** BOLD time course heritability magnitudes and spatial patterns are consistent in smaller subsamples. Scatter plots show that average BOLD time course heritability magnitudes (left) and spatial patterns in smaller subsamples of our data are consistent with those observed in the full sample (shading = standard deviation across 100 random subsamples at each percentage).

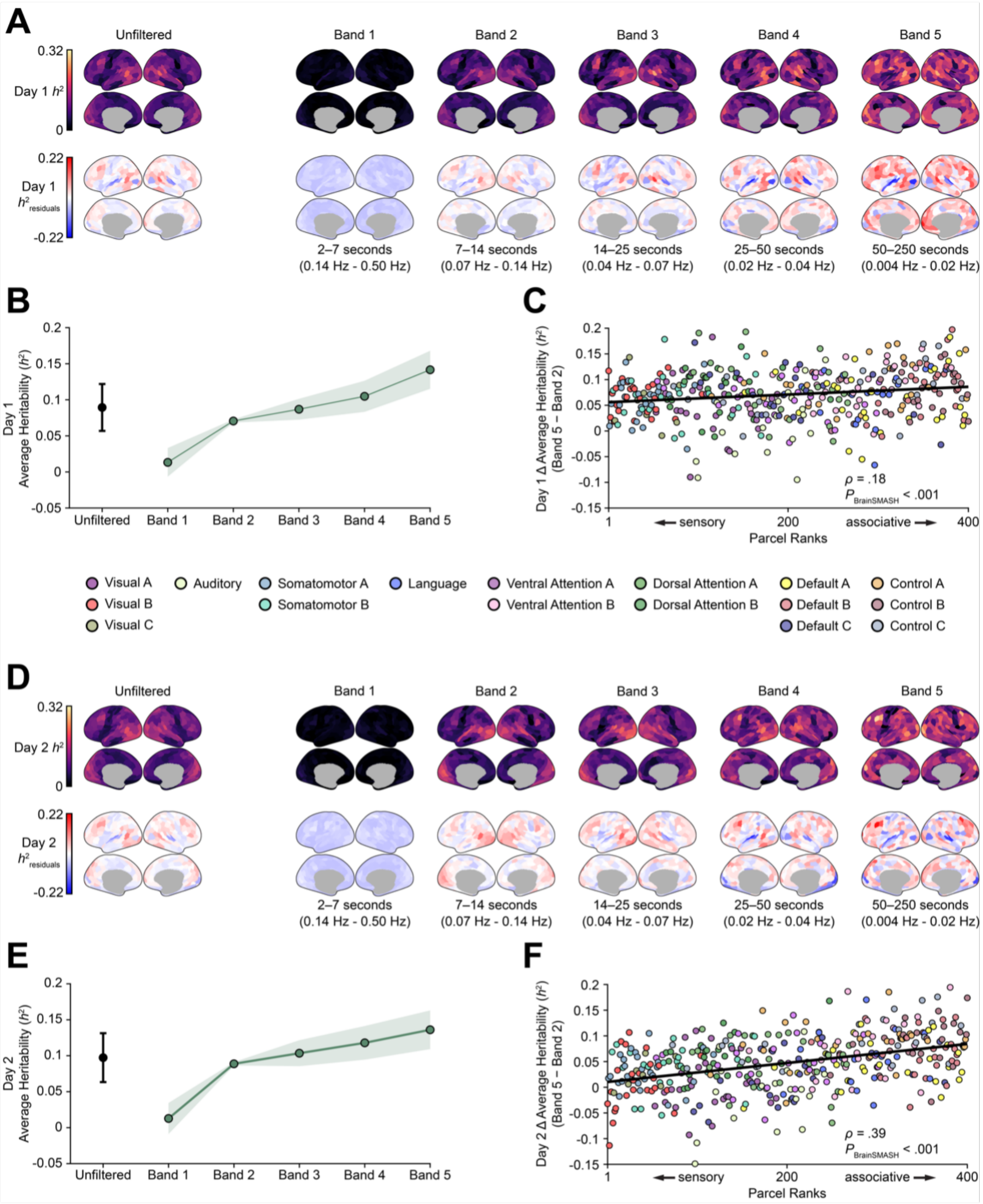


**Figure S6.** BOLD time course heritability is greater in slower frequency bands, especially for more associative parcels (for uncensored data). Same as Fig. 3 using full time courses for each day (including 20s rest and clip onset blocks).

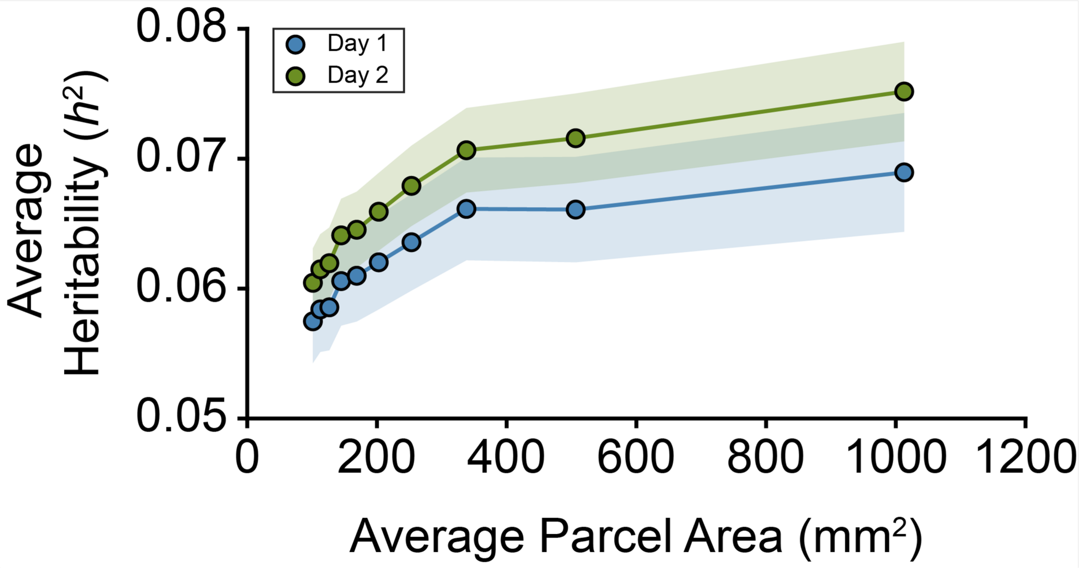


**Figure S7.** BOLD time course heritability is resolution-dependent. Scatter plots show that average BOLD time course heritability across all parcels is higher for coarser Schaefer parcellation resolutions (shading = SEM).

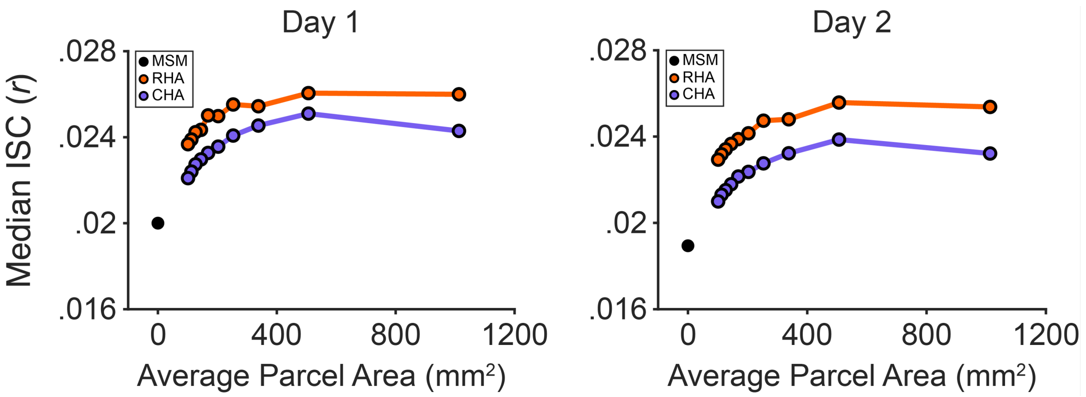


**Figure S8.** Hyperalignment increases ISC. Scatter plots show that, as expected, hyperalignment increases ISC (quantified as the median ISC value across all subject pairs and vertices), and that this increase is greater for response (vs. connectivity) hyperalignment and for coarser (vs. finer-grained parcellation resolutions).

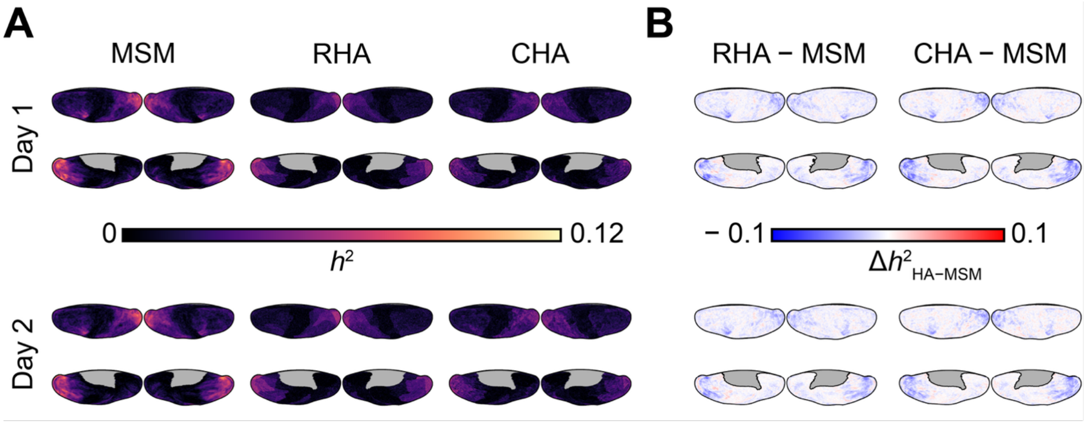


**Figure S9.** Hyperalignment reduces BOLD time course heritability. Dorsal and ventral views of the surfaces in Fig. 4C-D

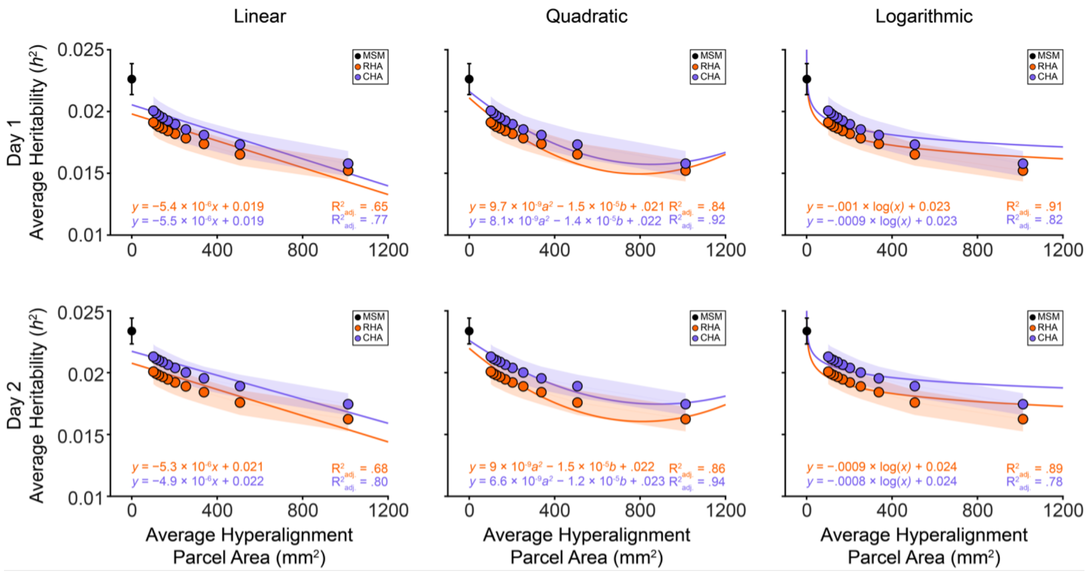


**Figure S10.** Linear, quadratic, and logarithmic fits of average BOLD time course heritability and hyperalignment resolution. Same as Fig. 4E for non-power law models of the relationship between hyperalignment resolution and heritability.

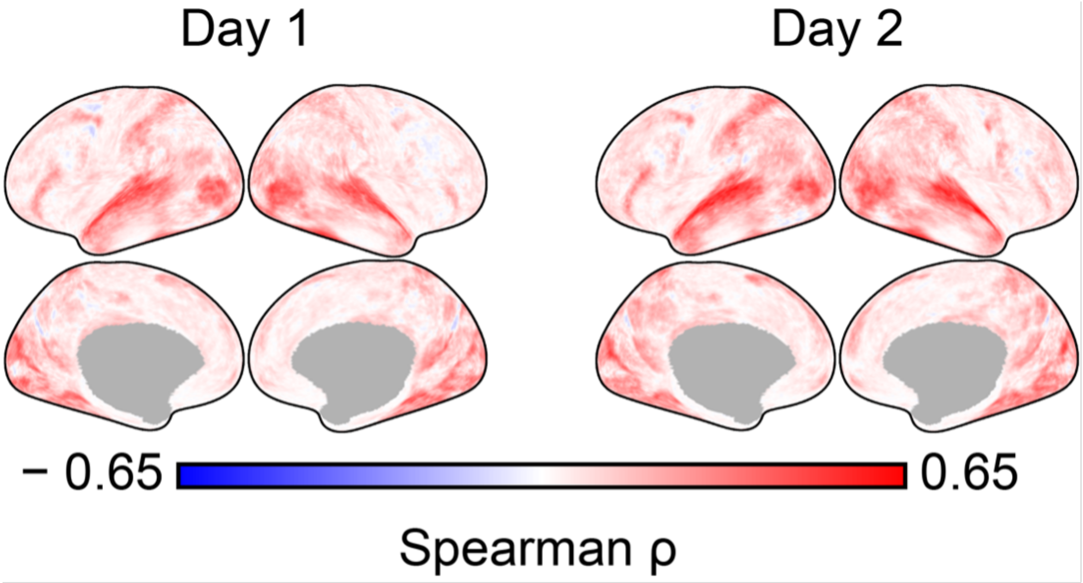


**Figure S11.** Subject pairs with longer NTs have more correlated BOLD time courses. Surface plots show that pairwise ISC values from one day of data collection scale with summed NTs from the other day's data, especially in auditory and visual cortices (n = 15,753 unique dyads; 54% of cortical vertices significant at FDR-corrected $P_{perm} < .05$ on both days).

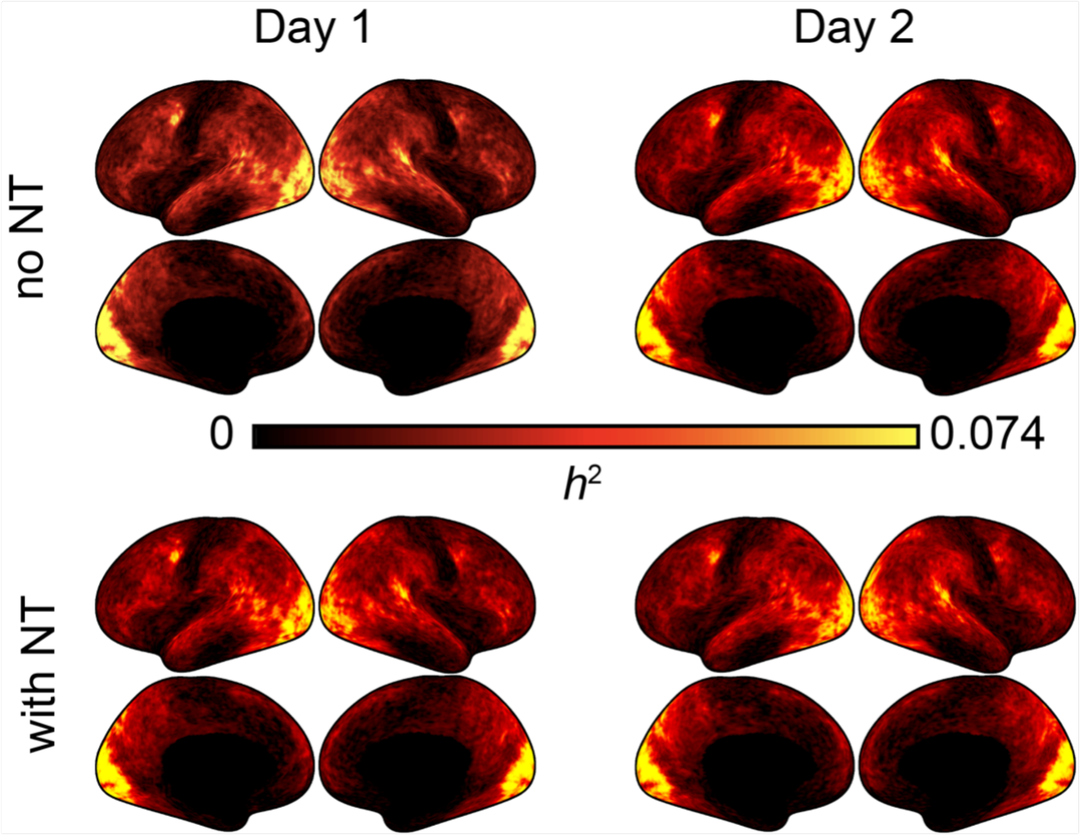


**Figure S12.** BOLD time course heritability with (bottom) and without (top) controlling for NT. Surface plots show the heritability maps used to generate the difference maps in the top row of Fig. 5B.

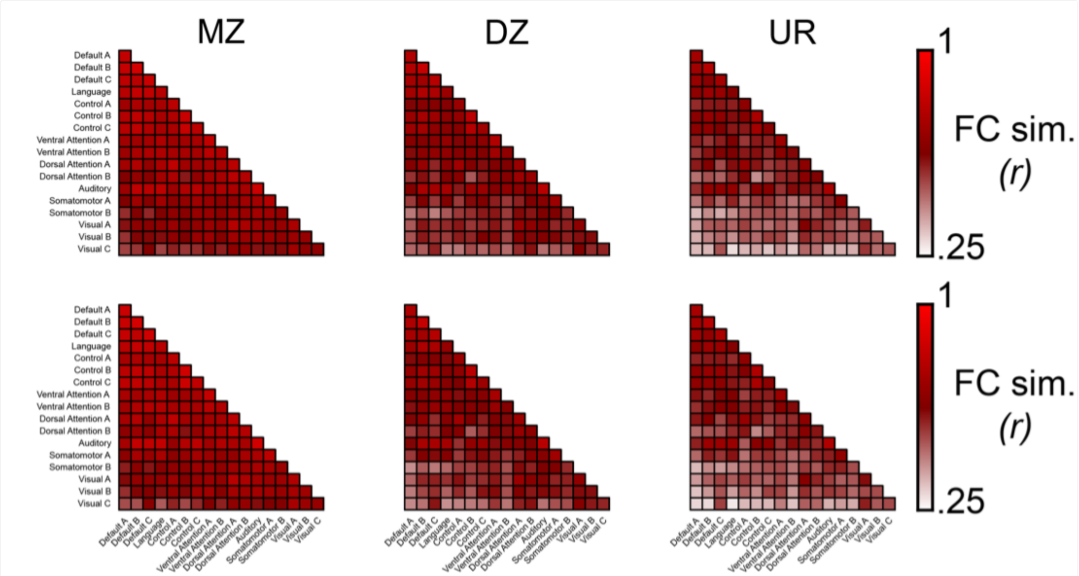


**Figure S13.** Movie-watching FC profile similarity by group. Same as S1 for FC profile similarity during movie-watching.

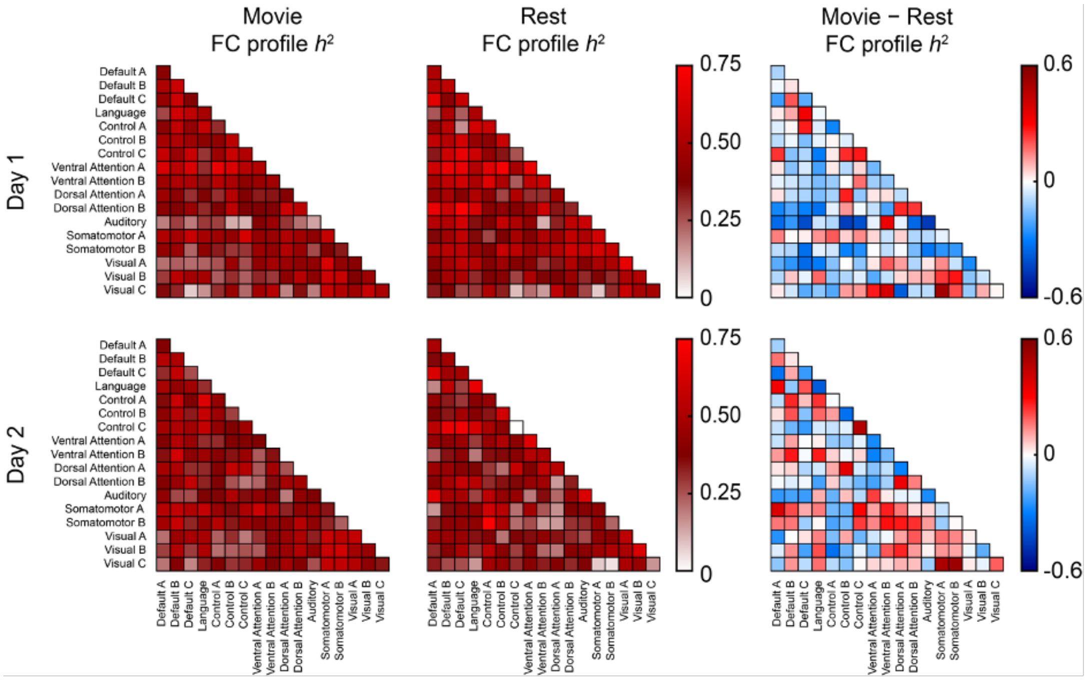


**Figure S14.** FC strengths are similarly heritable during movie-watching and resting states. Same as Fig. 3 but for FC strength (vs. profile) heritability. FC strengths were largely heritable during movie-watching (Day 1 mean $h^2$-SOLAR= .42 ± .13, Day 2 mean $h^2$-SOLAR = .41 ± .11; 84% of network combinations significant on both days at FDR-corrected $P_{perm} < .05$), but the cross-day reliability of the FC heritability patterns across network combinations was about half that of the FC profile analysis (Spearman ρ = .39, $P_{perm} < .001$), and no movie FC strength heritability values were significantly greater than rest FC values on both days for any network combination (right column).

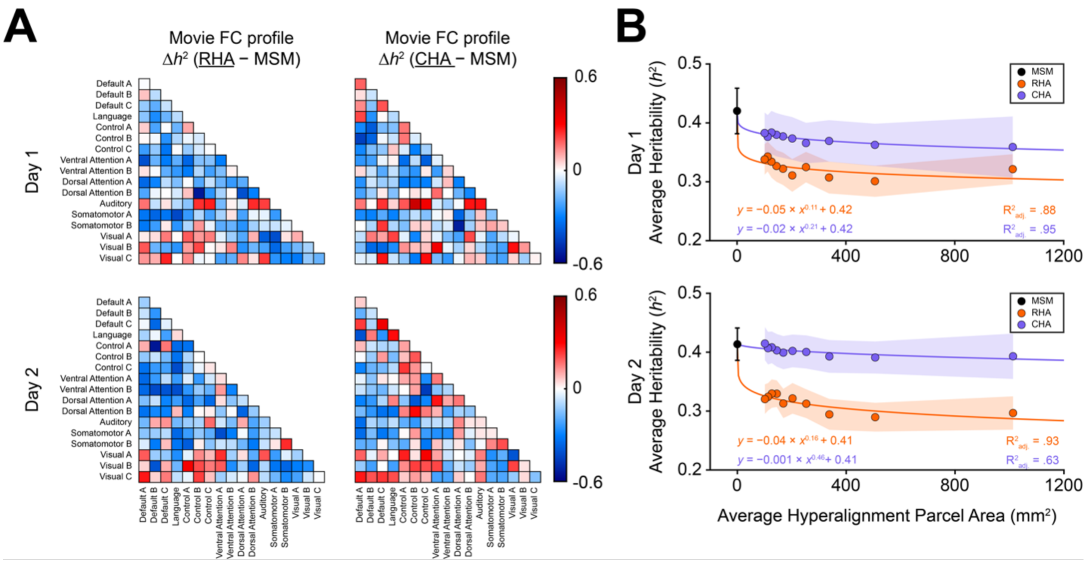


**Figure S15.** Response (but not connectivity) hyperalignment decreases FC strength heritability. (A–B) Same as Fig. 5 but for FC strength (vs. profile) heritability. RHA using the Schaefer 100 atlas decreased average FC strength heritability across all network combinations by 24% (95% CI: 12–35%) on Day 1 and by 28% (15–42%) on Day 2. Although CHA at the 100-parcel resolution lowered FC strength heritability on both days, these decreases were not statistically significant (Day 1: 15% [-10–39%], Day 2: 10% [-13–23%]).

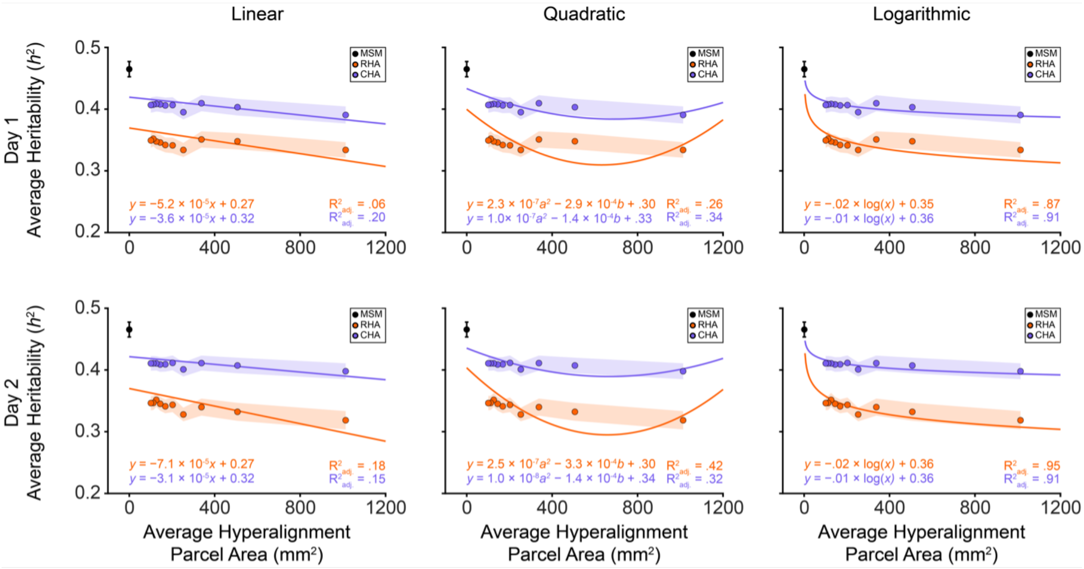


**Figure S16.** Linear, quadratic, and logarithmic fits of average FC profile heritability and hyperalignment resolution. Same as Fig. 4B for non-power law models of the relationship between hyperalignment resolution and FC profile heritability.

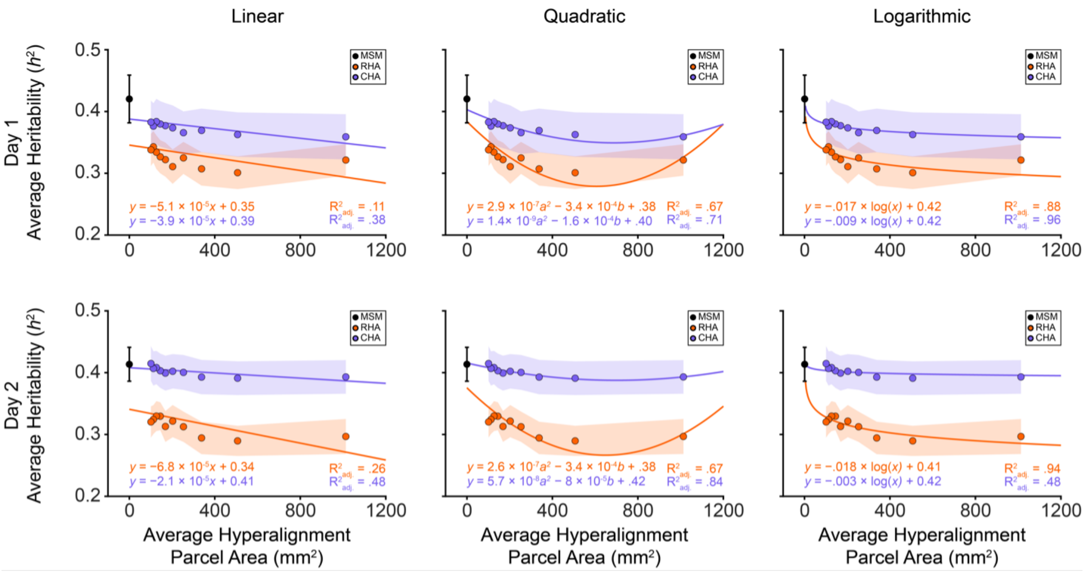


**Figure S17.** Linear, quadratic, and logarithmic fits of average FC strength heritability and hyperalignment resolution. Same as Fig. 4B for non-power law models of the relationship between hyperalignment resolution and FC strength heritability.
